# A real-time, multi-animal model for automatic face detection and identification of freely moving common marmosets based on YOLOv8 algorithms

**DOI:** 10.64898/2026.02.02.703330

**Authors:** Jiayue Yang, James Wang, Justine Cléry

**Author notes:** Corresponding author: Justine Cléry.

## Abstract

Precise and up-to-date information about animal location and identity allows us to better quantify individual behaviors in studies of neural activity, cognition, and animal health. In socially housed laboratory animals, identification is usually defined by observation or invasive markers, making the data collection time-consuming, variable across experimenters, and disruptive to animals. We established an automatic pipeline for real-time identification of common marmosets in captivity using a close-view camera. It uses the supervised deep-learning YOLOv8 model to localize individuals, detect faces, and classify identities. Moreover, we use recognition of uniquely color-coded collar beads to improve detection accuracy among visually similar individuals. Across adult and juvenile marmosets, our system automatically identifies marmosets with > 82.9% precision and > 91.5% recall, achieving human level performance. This pipeline is designed to be easy-to-use and generalizable across non-human primate species, ages, and recording hardware, providing rapid and automatic identity recognition from real-time video.

## Introduction

In behavioral neuroscience, an accurate identification of laboratory animals is crucial for welfare assessment and ethical care (National Research Council (U.S.), 2011; Vidal et al., 2021). More importantly, the animal identity is also required when conducting individual-specific behavioral experiments. However, identifying specific animals in a family group can be challenging, especially if they are not marked and share similar facial features. Invasive recognition methods, including branding, tattooing, and ear tagging, may disrupt animals’ behaviors and cause potential stress (Carstens and Moberg, 2000; Lim et al., 2019; Roughan and Sevenoaks, 2019). To avoid such harms, video analysis is becoming more popular as a non-invasive method to identify the animals by their appearance, with no direct contact or handling being involved (Norouzzadeh et al., 2018; Schindler and Steinhage, 2021). However, collecting video data of animals that is subsequently labelled by human observation can be time-consuming and hard to reproduce across experimenters. (Buchan et al., 2003; Marion et al., 2020). Therefore, considering replacement of human work, machine learning tools have been introduced and increasingly used, providing an accurate and automatic estimation of animal identities.

Computer vision and machine learning have made many advancements and offered the foundation to build these automatic systems (Sturman et al., 2020). Within these approaches, deep learning, a subset of machine learning, has been widely used in animal identification based on video and image data. Specifically, by utilizing deep learning with video recordings, object detection allows an accurate localization and identification of animals, even with those who live in complicated environments (Yu et al., 2018; Zhuang et al., 2025). Algorithms like ResNet (He et al., 2015) and You Only Look Once (YOLO) models (Jocher et al., 2023; Redmon et al., 2015) have demonstrated promising accuracy and computational speed in identifying animals in naturalistic environments (Bakana et al., 2024; Petso et al., 2021). In addition, unique facial features can also be used as inputs to classify and recognize animals. This approach has the advantage that it provides information regarding whether a specific animal is present or absent in the camera recording view at each timepoint. Although facial recognition has been demonstrated to be successful in wild and domestic animals (Bergman et al., 2024; Norouzzadeh et al., 2021; Schofield et al., 2023), its application in controlled laboratory settings is not yet extensively evaluated.

The common marmoset (*Callithrix jacchus*), a small non-human primate, is becoming more popular recently in neuroscience research for many reasons (Kishi et al., 2014; Okano, 2021; Sasaki et al., 2009). Marmosets are family-bonded, cooperative, and notably pro-social. Moreover, they can perform multiple series of behavioral and cognitive tasks, such as observational and reversal learning (Koski and Burkart, 2015; Miller et al., 2016). As marmosets are typically maintained in social family groups (Yoshimoto et al., 2018), accurate identification of specific individuals is therefore essential for assessing which animal is performing a given task, which also enables a tracking of their behavioral performance over time. However, existing identification techniques of marmosets usually involve wearable micro-sensing devices, requiring stable device positioning to the sensor and constant adjustments to ensure the devices fit the animal. This automatic system can also become unreliable during rapid movement or if multiple marmosets are present at the sensor, which limits the consistency and stability of identity tracking in freely moving marmosets during their tasks. Thus, facial recognition offers a non-invasive alternative, without a strict need of a physical identity marker, enabling a continuous, accurate, and automated identification when the marmoset enters the designated task performance space.

Here, we developed a pipeline for automatically detecting and identifying marmosets simultaneously from real-time videos, based on their faces. Previous studies have introduced the feasibility of the methods we used in this study (Dave et al., 2023; XiaoAn et al., 2024), of which we adapted them for facial-based marmoset recognition. We apply the YOLOv8 model (Jocher et al., 2023), trained on images of specific individuals (three adult and two young marmosets), to generate a real-time marmoset recognition on a frame-by-frame manner. In addition to marmoset faces, we use the color-coded bead on individual collars to facilitate the identity prediction accuracy. Finally, we further evaluated our model performance on a dataset from both adult marmosets and young marmosets at different developmental stages. We show that our facial-based pipeline achieves high detection performance comparable to human expert level, with a strong generalizability across recording settings and marmosets at various ages.

## Methods and Materials

### Animals

Three adult common marmosets (*Callithrix jacchus*, 1 father, 1 mother, 1 son, 6 years weighted at 464g, 5 years at 566g, and 3 years at 450g respectively) and two young common marmosets (two female twins, average weights 329g and 338g, aged 7 – 11 months) were involved in this study. The three adult marmosets belonged to the same family unit (parents: Adult1, Adult2, and their adult offspring: Adult3), while the young marmosets were twin siblings (Young1 and Young2) from another family. The marmosets are housed at The Neuro’s animal facility in family units in indoor enclosures. Housing cages included two sizes. The first size housed three adult marmosets with dimensions of 1.372m length x 0.760m width x 2.092m height, and the second cage that housed the two young marmosets with their family has the dimensions of 1.065m length x 1.067m width x 2.092m height). We placed a collar with a uniquely colored bead on each marmoset to allow consistent visual identification during the experiment. All experimental procedures were conducted under the Canadian Council on Animal Care guidelines, Standard Operating Procedures of marmosets, and Animal Use Protocol (AUP# 10000 and 10001) approved by The Neuro and McGill’s Animal Care Committee.

### Video collection

Marmoset face dataset was collected by mounting a primate testing chair with an added camera component to the door of marmoset housing cages (Figure 1A and 1C). High-resolution videos were acquired using an industrial color/RGB camera (JIERUIWEITONG DF200-1080P). The industrial camera was selected specifically for its small size (camera box: 36mm width x 36mm height) and capability of close-distance recordings. High-resolution recordings (1920×1080 pixels, 30 frames per second) obtained with the small focal length lens (2.8mm) enabled a wide field of camera view. This allowed the collection of full facial features and posture variability at close recording distances. The camera was adjusted for focus and placed approximately 10.50 cm from the housing cage door (Figure 1A and 1B). The camera was fixed in position using a transparent protective case, which was designed to prevent damage to the camera by the marmosets (Figure 1B).

**Figure 1.**
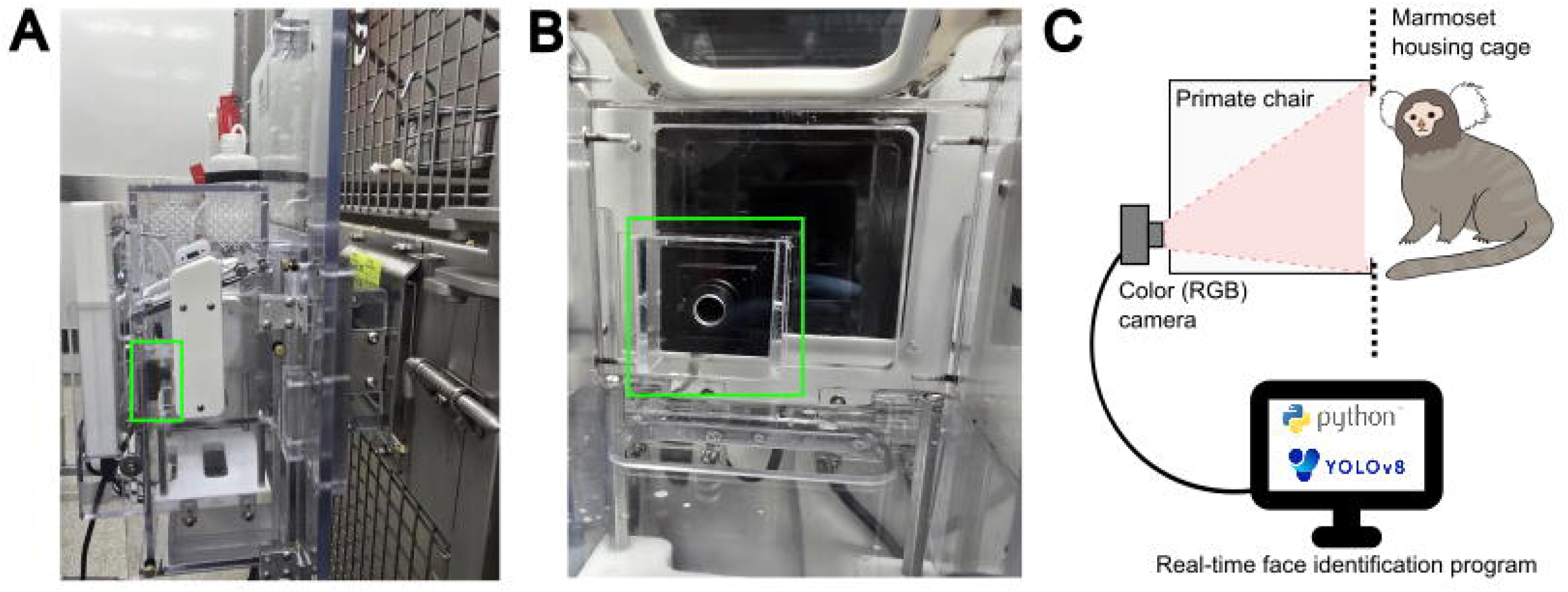
Experimental setup of the camera system and primate chair for face detection and identification. (A) Side view of the camera (green box) and primate chair, attached to the housing cage. (B) Frontal view of the camera fixed on the primate chair, enclosed within a protective cover. (C) A schematic illustration of the real-time facial image recording and automatic identification of the marmoset entering the primate chair.

Prior to the experiments, marmosets were acclimated to entering the primate testing chair space. This space minimized the chair surface reflection, reducing interference between true facial features and reflections during face detection and identification. Three adult marmosets from one family were recorded for 1 hour across each of the 5 recording days, with unrestricted voluntary access to the primate chair space. For the adult marmosets, the housing cage door was opened at the beginning of the recording session allowing them to enter and exit freely into the primate chair space for food rewards and observation (Figure 1C). Two young marmosets were briefly isolated and recorded separately for testing and improving the automatic face extraction program. We recorded them at two developmental timepoints, 7 months old and 11 months old (an additional time point at 16 months old has been added for one marmoset to test the identification without collar). During video collection, a sliding panel and an in-cage box were positioned near the housing cage door to temporarily isolate individual marmosets from other family members. Individual isolation was kept brief (approximately 10 minutes) to prevent disturbance and potential stress due to family separation. Marmosets accessed the primate chair space through the housing cage door (11cm width x 10.16cm height).

### Video preprocessing and bounding box annotation

The collected videos were preprocessed to identify which video clips included valid face and collar information of the marmoset. We defined valid face and collar features by whether the full-face details and colored bead appeared visible from the clips (Figure 2A, Step 1). To capture sufficient variability in postures, individuals, and lighting conditions, clips were sampled throughout the adult marmoset videos (approximately 5 hours) (Figure 2B). We used OpenCV to extract frames from the selected clips, and the extracted frames were reviewed to exclude those that were blurry (Bradski, 2000). Computer Vision Annotation Tool (CVAT) was applied to perform bounding box annotation of marmoset faces and collar colors from the extracted frames (Sekachev et al., 2020).

**Figure 2.**
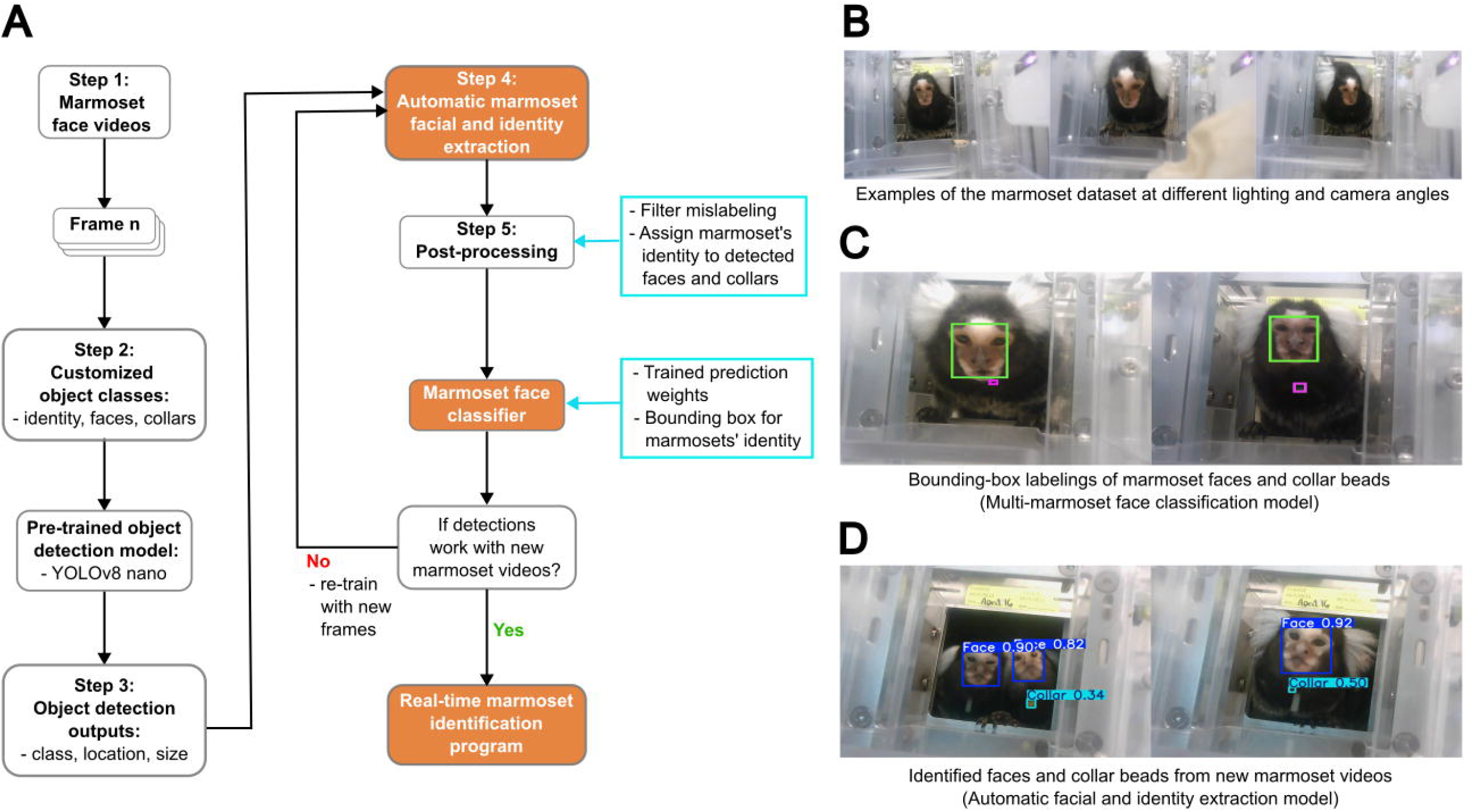
Workflow and design of the marmoset facial detection and identification model. (A) The architecture of the real-time marmoset facial recognition program. (B) Marmoset face images from three camera angles. (C) Bounding boxes of marmoset faces (green box) and collars (pink box) were manually labeled in the training and validation datasets to train the multi-marmoset face classification model. (D) The schematic of the automatic face (blue box) and collar bead (cyan box) extraction model.

### Marmoset face and identity dataset

To minimize image computations and data storage, we created a dataset of 2498 annotated images from the three adult marmosets. All images were manually annotated to label marmoset faces, individual identities, and the collar bead colors (Figure 2C). The annotated images were used for training models of multi-marmoset face classification and the automatic identity extraction, which can automatically detect, localize, and identify marmoset faces (Figure 2A, Step 1 – 4). We created another dataset of two young marmosets at 7 months old (total images = 502) for testing the automatic facial and identity extraction (Figure 2A, Step 4 – 5). For both adult and young marmoset datasets, images were randomly divided into a training set and a validation set at a ratio of 8:2. Moreover, we evaluated the model performance and its generalization across developmental stages using new videos: 1) from the adult marmosets and 2) from the same young marmosets at 11 months, which were not involved during initial program training.

### Marmoset face recognition pipeline

The marmoset facial detection and identification was primarily based on the YOLOv8 algorithm (Jocher et al., 2023; Varghese and M., 2024). The architecture of the program consists of two main models: a multi-marmoset face classifier and an automatic facial and identity extraction (Figure 2A). In the initial step, we deployed multiple pre-trained YOLOv8 models (YOLOv8 nano, YOLOv8 small, and YOLOv8 medium) on adult marmosets’ dataset (total images = 2498) to train the multi-marmoset face classifier (Lin et al., 2015). Training performance and detection accuracy of these models, generated by YOLO metrics as a CSV file, were evaluated to select the optimal model for subsequent training (see Results section). These pre-trained models share the same object detection backbone but differ in number of parameters and computing power. Larger models, such as YOLOv8 medium, provide higher detection accuracy but require greater computational resources and longer inference time. In contrast, smaller models prioritize the computational efficiency. The YOLOv8 nano model was selected and used for the training of all models in the program (Figure 2C). The adult marmosets’ dataset was also applied in the training of automatic facial and identity extraction. This trained automatic face extraction was utilized on the young marmoset dataset (total images = 502). Since young marmosets were recorded individually, we assigned the automatic annotation of marmoset faces and collar beads based on their recorded identity (Figure 2D). All automatic labeling was manually reviewed to remove mislabeled annotations. We used this dataset (total images = 449) to train the multi-marmoset face classifier for the young marmosets.

The marmoset facial recognition program was trained in Anaconda virtual environment. The model training terminated at the early stopping criteria, when no improvement was observed across 100 most recent epochs (i.e. training iterations). This tool involved training of three YOLOv8-based programs: (a) multi-marmoset face classifier for three adult marmosets was trained for 183 epochs; (b) automatic facial and identity extraction was trained for 124 epochs; (c) multi-marmoset face classifier for two young marmosets was trained for 245 epochs. The final models generated 2D bounding boxes, assigning identity and collar information of individual marmosets. The detection results of input videos were exported as text files, which represented the corresponding label identity, the bounding box size, and its location (Jocher et al., 2023; Redmon et al., 2015).

### Model evaluation and analysis

The marmoset facial recognition program was evaluated using a range of performance evaluation metrics. The evaluation metrics included: precision, recall, Mean Average Precision (mAP), Validation Distribution Focal Loss (DFL), and training time. We also computed the F1 score to select the best setup to be applied in the final real-time detection pipeline. In addition, we calculated the inter-individual face similarity between marmosets to test whether the biological face similarity between family members can explain the potential mislabeling of the face classifier programs.

#### Intersection over Union (IoU)

Intersection over Union (IoU) calculates the amount of spatial overlapping between a predicted bounding box and the ground truth (i.e. manually labeled bounding box) (Jocher et al., 2023; Rezatofighi et al., 2019). It measures the localization accuracy and error amount between the predicted annotation and the manually labeled ground truth. A detection was considered correct if the IoU reached specific thresholds. In this study, we reported the mean averaged precision (mAP) at the IoU thresholds from 0.5 to 0.95 (increments = 0.05). The IoU is calculated by the area of overlap (*A* ⋂ *B*) and area of union (*A* ⋃ *B*) of the predicted bounding box (A) and ground truth bounding box (B). If IoU = 1, the predicted bounding box is perfectly aligned with the true annotation. If IoU = 0, there is no overlap between the two boxes.

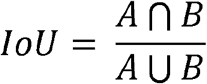

#### Precision, recall, and F1 score

Precision, recall, and F1 score are commonly used in the evaluation of classification models (Dehmer and Basak, 2012; Powers, 2008). Precision measures the overall proportion of correct face identifications over all positive results, indicating the accuracy of the detection model. Recall quantifies the proportion of the face identification that are correct compared with all actual positives, which reflects the ability of the model to predict correct marmoset face and collar beads. The F1 score is the harmonic mean of precision and recall, considering the false positives and false negatives in the model assessment.

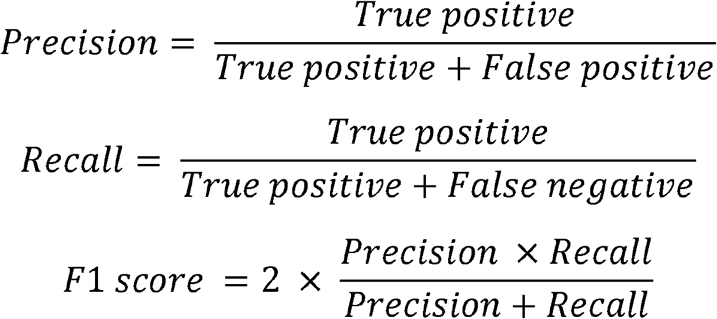

#### Mean Average Precision (mAP)

Mean Average Precision (mAP) assesses how accurate the model detects objects and how well the model localizes the object in the image. In object detection algorithms, precision is the number of correctly detected objects, with recall being the number of true objects that are successfully detected. As the precision and recall are affected by different detection confidence thresholds, the performance is evaluated by the precision-recall curve, which plots the model precision (y-axis) against recall (x-axis) for a particular threshold. The average precision (AP) is the area under the precision-recall curve (Padilla et al., 2021). Before calculating the AP, the precision-recall pairs are interpolated to create a monotonically non-increasing precision-recall function, meaning that the interpolated precision (*P*_*interpolation*_) is the maximum precision (*max*_*j*≥*i*_ *precision*_*j*_)t the recall level larger than or equal to Recall_i_. This interpolation assigns a highest precision value at each recall level, which smooths the precision-recall curve and reduces the impact of measurement fluctuations and noises to the evaluation of model performance.

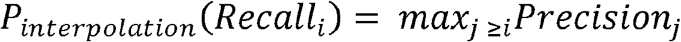

For each label class in the object detection model, the AP is the area under the curve (AUC) of the interpolated precision-recall curve, calculated by summing the interpolated precision weighted by the incremental increase in the recall (Maxwell et al., 2021; Padilla et al., 2021).

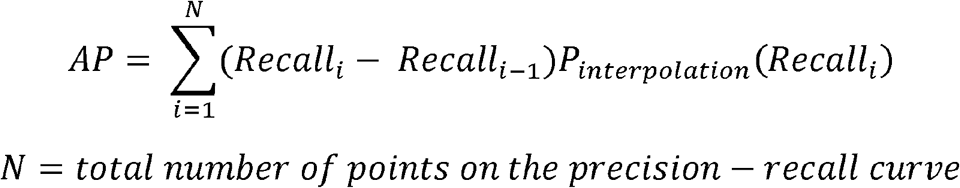

The AP is extended further to calculate the Mean Average Precision (mAP), which is the averaged AP values of different label classes in the object detection model (total class number = n). The APi is the mean AP calculated per label class at index *i* at multiple IoU levels (IoU = {0.50, 0.55, …, 0.95}). This calculation generates a more comprehensive evaluation with multiple label classes and their predicted localization accuracy of our marmoset facial recognition program.

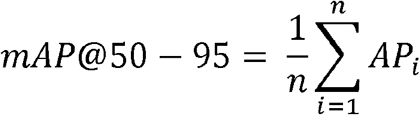

#### Validation Distribution Focal Loss (DFL) and training time

The distribution focal loss (DFL) and training time were also used to support the selection of the final marmoset facial recognition model. The DFL function measures the model’s performance to refine the bounding box predictions, based on the localization uncertainty of the bounding box annotations (Jocher et al., 2023). The predicted bounding box position is compared with the ground truth annotations, with a lower DFL value indicating a more precise bounding box prediction.

Training time was used the training time to assess the efficiency and the feasibility of the real-time marmoset face recognition program. The training time estimated the computational cost of the program, which provided additional information to select models with similar accuracy. The training time is calculated by the total time duration to train the model. For the number of training epochs, N_epoch_, the total training time is calculated through multiplying N_epoch_ by the training time per epoch (t_epoch_):

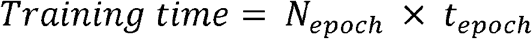

#### Inter-individual face similarity analysis

We calculated the inter-individual face similarity using customized Python scripts (See Data Availability). The image feature embeddings, based on YOLOv8 algorithms, calculate the numerical representations (i.e. vectors) using a computer vision model, which encodes the semantic and visual maps of the image (Long Chai et al., 2023; Varghese and M., 2024). In our marmoset facial recognition program, the intermediate layers included extensive feature maps of the visual details of the marmoset face images. By extracting and analyzing the image feature embeddings, we obtained various numerical representations of the marmoset faces, which allowed us to compare face shapes, distances between face features (eyes, mouth, nose, etc.), and the fur patterns of different individuals. We assessed the inter-individual face similarities for 2 family groups: (a) within the adult marmoset family (parents and adult offspring) and (b) between the two young marmosets (twin siblings). For each individual marmoset, we selected 10 images with similar head orientation, lighting conditions, and camera angles, and cropped them to only include the face regions. As the marmoset face images passed through the YOLO convolutional layers, the images were outputted into a feature map (Aghdam and Heravi, 2017; Dumoulin and Visin, 2016; Long Chai et al., 2023; Redmon et al., 2015). For one marmoset face image, the feature map (F_i_) is encoded in the number of inputs (n_input_) and the feature map shapes (Aghdam and Heravi, 2017; Dumoulin and Visin, 2016).

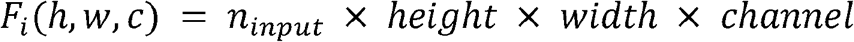

We extracted face embeddings from the face classifiers of the adult marmosets’ family (parents and son) and young twin marmosets from another family. To generate a single vector (i.e. embedding, e_i_) representing the marmoset face feature per channel, we averaged across the spatial dimension (i.e. height and width of the image) per channel for one marmoset face image (Lee et al., 2025). The extracted embedding (e_i_) was averaged across the 10 images for each individual marmoset, which allowed us to have an averaged embedding vector (e_marmoset_) and feature maps to calculate the inter-individual face similarities. We preformed L2 normalization on this embedding (e_marmoset_) to create a unit vector (e_norm_) for inter-individual comparison, using scikit-learn preprocessing function (Pedregosa et al., 2011).

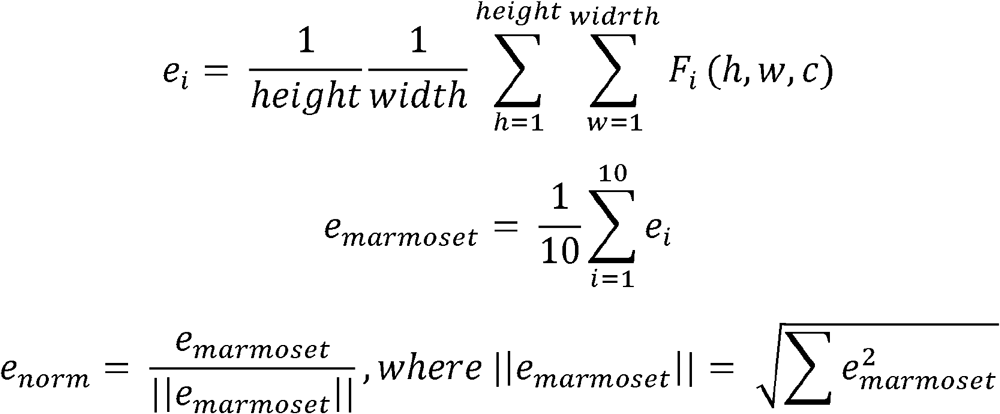

As such, with the normalized embedding vectors from the marmosets in this study, we calculated the inter-individual face similarities between marmoset pairs using cosine similarity (Li and Han, 2013; Nguyen and Bai, 2011; Thongtan and Phienthrakul, 2019). Moreover, we also calculated the Euclidean distances between extracted embeddings from two marmosets (Jozwik et al., 2022; Sugase-Miyamoto et al., 2014). The calculation equations were described using the normalized embedding vectors of a marmoset pair (e_norm – 1_ and _enorm - 2_) in scikit learn library (Merchant et al., 2023; Pedregosa et al., 2011):

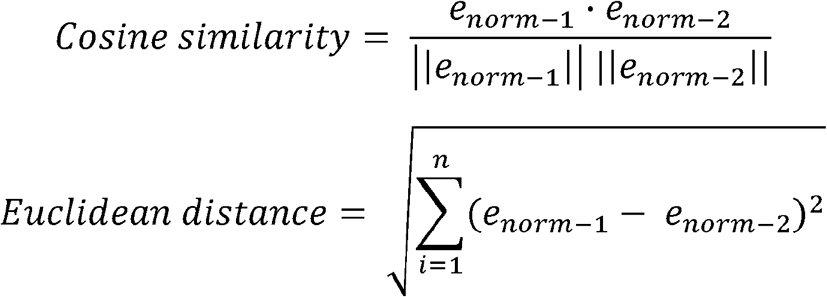

We plotted a violin plot of cosine similarity and Euclidean distance to interpret the inter-individual facial similarities calculated from the marmoset pairs’ face embeddings. In addition, the similarity values were standardized using within-model z-scores.

Statistical tests were performed only within the face classifier of each marmoset family, as embedding spaces may vary in scaling, learned features, and baseline metrics making cross-model comparison of inter-individual face similarity unreliable (Bollegala, 2017). To test whether the learned facial embeddings between different marmosets is related to the biological facial feature similarities, we performed the pairwise Welch’s t-test and the Cohen’s d to examine the significance and effect size. Within each family unit, facial similarity was compared between two types of family relationships (e.g. mother-son vs. father-son).

### Real-time face recognition program

We performed the real-time marmoset face detection and identification using the customized Python scripts, based on the trained weights from the final multi-marmoset facial classification model in YOLOv8. Live videos were collected using the color (RGB) camera and simultaneously processed frame-by-frame to detect the marmoset identities. For each detected bounding box, the scripts returned a corresponding label of marmoset face and collar bead color. We assigned the detected collar beads as the corresponding marmoset identity with a higher weight, which improved the detection confidence across frames.

To ensure the program accuracy and efficiency, we temporally stored the detection results of most recent 30 frames (approximately one second). The most frequently detected marmoset identity within this time window was exported as the current output. We wrote and updated the detection results continuously into an output JSON file, enabling real-time identity reading of the marmoset who entered the primate chair most recently.

## Results

### Comparison of pre-trained YOLOv8 models

Model performance included the precision (the proportion of correct positive predictions), recall (the proportion of corrected predicted ground-truth labels), and mAP@50–95 (the average detection accuracy across different IoU object localization thresholds; *see Methods and Materials section for detailed definitions*). The results of the training and evaluation of the pre-trained YOLOv8 models (YOLOv8 nano, YOLOv8 small, YOLOv8 medium) were shown in Table 1. The models reached their best performance at 180 epochs (YOLOv8 nano), 132 epochs (YOLOv8 small), and 124 epochs (YOLOv8 medium). Each of the models was trained until reaching the early stopping criteria (i.e. no improvement within the last 100 training epochs). By comparing the performance differences between the YOLOv8 pre-trained networks, we aimed to find the most suitable model for the real-time marmoset identification program.

**Table 1.** Performance comparison of multi-marmoset face classification models. Comparison of Recall, Precision, F1 score, mAP at IoU = 0.5:0.95, Validation DFL, and training time of the three pre-trained model, based on the performance of marmoset face detection and identification on the adult marmosets’ dataset. The highest values of each parameter were highlighted in bold.

| Metrics | YOLOv8 nano | YOLOv8 small | YOLOv8 medium |
| --- | --- | --- | --- |
| Recall | **0.964** | 0.957 | 0.956 |
| Precision | 0.932 | 0.927 | **0.937** |
| F1 score | **0.948** | 0.942 | 0.946 |
| mAP@50-95 | 0.710 | 0.710 | **0.713** |
| Val DFL | **0.917** | 0.953 | 1.010 |
| Training time (s) | **5544** | 5908 | 10705 |

As presented in Table 1, YOLOv8 nano model reached the best overall detection performance of the adult marmosets’ dataset. YOLOv8 medium model achieved a higher precision among all three pre-trained networks. However, the recall and F1 score were higher for the YOLOv8 nano model, indicating robust model sensitivity and a balanced precision-recall value. Mean average precision (mAP@50–95) showed values ranging from 0.710 – 0.713 for the three models. YOLOv8 medium model reached the highest detection accuracy for adult marmoset faces, though the difference was minor while comparing to the YOLOv8 nano and small models. In addition, we considered the validation DFL and training time to support the selection of the optimal pre-trained model. Validation DFL increased with model sizes, suggesting that the detection localization became less accurate from YOLOv8 nano to YOLOv8 medium. Moreover, the training time increased from YOLOv8 nano to medium model, while YOLOv8 medium took about twice the training time compared to YOLOv8 nano model. Thus, we selected the YOLOv8 nano model for training the real-time program due to its high computational speed and efficiency, while maintaining reliable marmoset face prediction compared to larger YOLOv8 models (Jocher et al., 2023).

### Prediction based on adult marmoset faces shows reliable identification

We applied the YOLOv8 nano model for the adult marmoset face classification. The training curves of the overall precision, recall, and mAP@50-95 (IoU = 0.5:0.95) were visualized in Figure 3A, B, C, and Figure 3—figure supplement 1. The face classifier weights were selected at the training epoch 183 (denoted by the red dotted line), reaching the precision at 0.932, recall at 0.964, and the mAP@50-95 (IoU = 0.5:0.95) at 0.710 (see Table 1). We evaluated the face classifier performance per label class using the adult marmoset’ validation dataset (Figure 3D, see Appendix A—Table 1). Analyzing the label classes, the adult marmosets face classifier showed stable and high precision and recall across both individual identity (i.e. Adult1, Adult2, Adult3) and bead-color (i.e. collar_Adult1, collar_Adult2, collar_Adult3) label classes (Video 1). In addition, the marmoset face classes showed high mean average precision across varied IoU thresholds, suggesting a robust prediction localization for marmoset faces (mAP@50-95 = 0.841, 0.787, 0.868 for three adults respectively). The lower mAP value observed for the bead-color label category (mAP@50-95 = 0.603, 0.665, 0.515 for collars of three adults) suggested a greater variability in bounding box localization for the marmoset collar beads. This effect was consistent with the small bounding box sizes of the collar-bead label class, which increased the model sensitivity to localization variability in the prediction (example detection shown in Figure 2C). The normalized confusion matrix comparison was shown in Figure 4. The detection accuracy showing similar performance using the adult marmosets’ training dataset (Figure 4A) and validation dataset (Figure 4B), with very low occurrence of false detection between the marmoset individuals and the background images.

**Figure 3.**
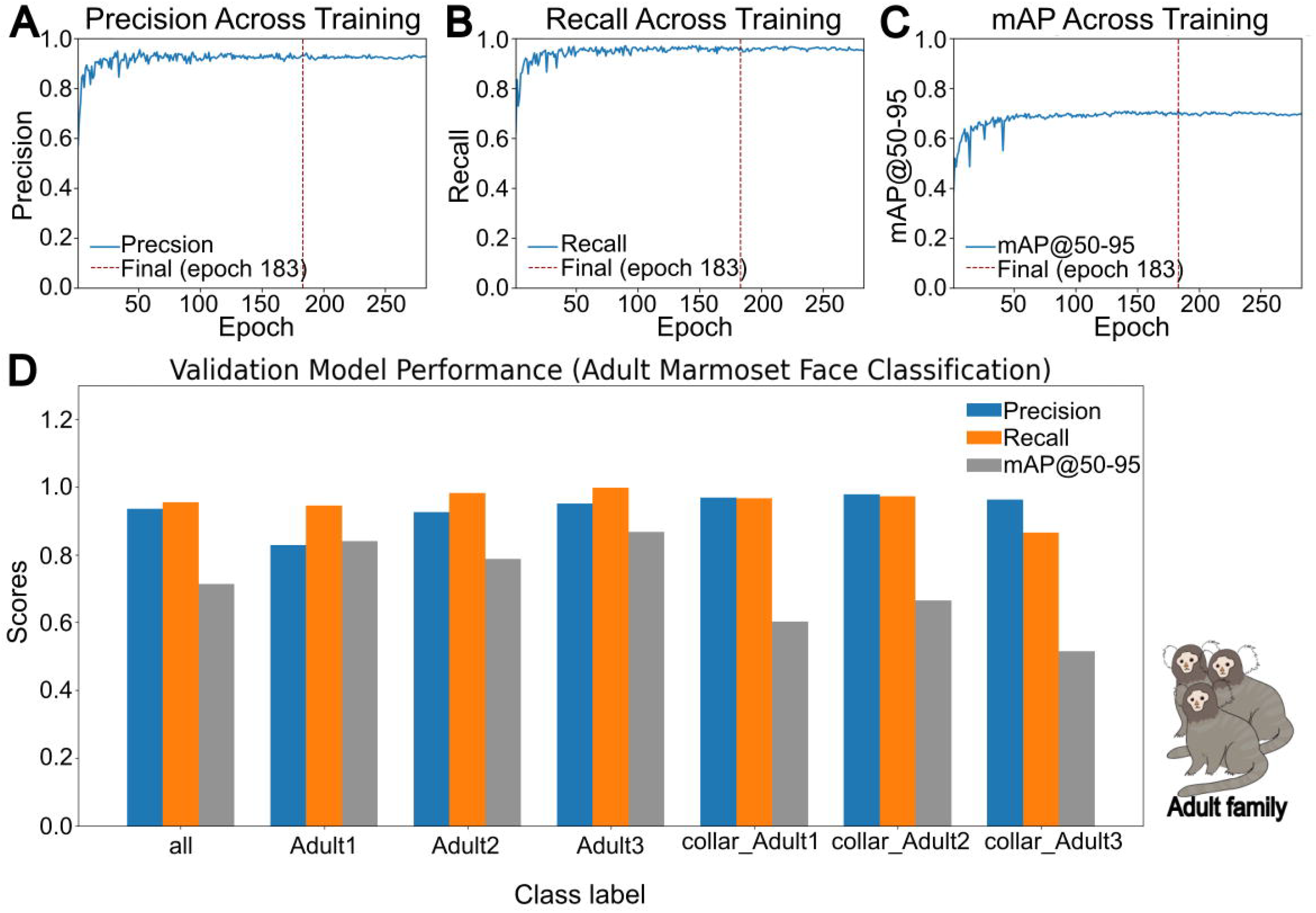
Training performance of the multi-marmoset face classification model for three adult marmosets, using the pre-trained YOLOv8 nano model. (A) Precision of all detection classes across training epochs. The red dotted line denoted the final model with the best performance at training epoch 183. (B) Similar to (A), except for overall recall. (C) Similar to (A), except for the mAP at the IoU at 0.5:0.95. (D) Overall Model precision, recall, and the mAP@50-95 (IoU = 0.5:0.95) for each label class.

**Figure 4.**
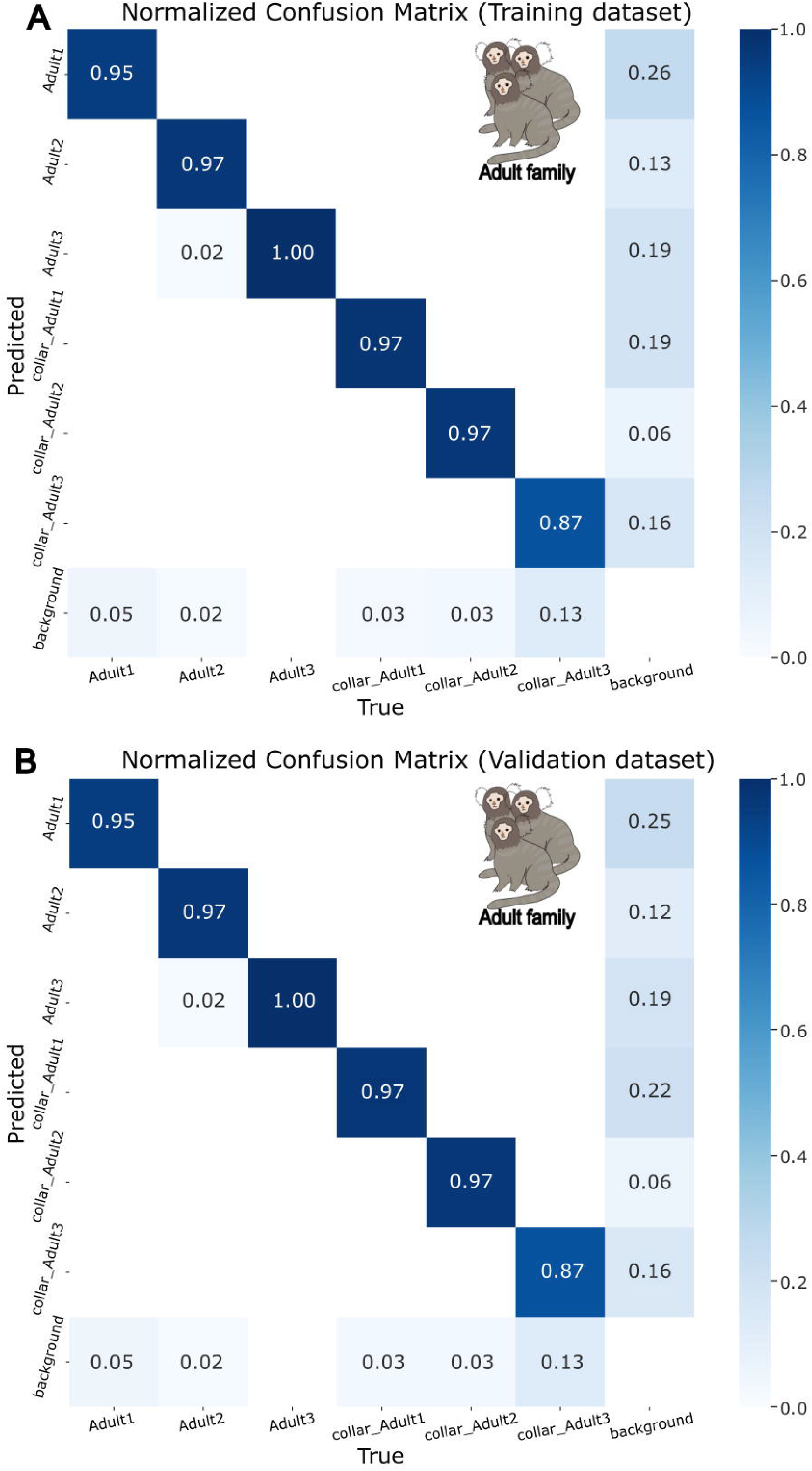
Normalized confusion matrix per-class classification across the 6 label classes in the adult marmoset recognition model. The y-axis represents the predicted class, and the x-axis represents the manually labelled class. Proportion was generated by the (A) training dataset and the (B) validation dataset, showing whether certain classes were frequently mislabelled as a different class.

### Automatic facial and identity extraction accurately localizes new marmoset faces and their collar beads

The labelled adult marmosets’ dataset was used for training the automatic facial and identity extraction, using the YOLOv8 nano pre-trained model. This framework aimed to extract and localize the marmoset faces and collar beads from the collected video, including new marmosets. The trend of the training was shown in Figure 5A – C and Figure 5—figure supplement 1, where the detection accuracy improved rapidly in the beginning, with the increasing rate then stabilizing and reaching a plateau. We selected the final weights at the training epoch of 124, with a prediction score at 0.940, a recall score at 0.970, and the mAP@50-95 (IoU = 0.5:0.95) at 0.716. In addition, precision, recall, and mAP@50-95 (IoU = 0.5:0.95) were calculated per label class for marmoset faces and collar beads (Figure 5D). Based on the model performance of the adult marmosets’ validation dataset, the final face extraction system accurately predicted the marmoset faces (precision = 0.919, recall = 0.968) and collar beads (precision = 0.957, recall = 0.95). The bead-color class showed lower mAP@50-95 (IoU = 0.5:0.95) values at 0.6, compared to the mAP@50-95 (IoU = 0.5:0.95) for marmoset face class at 0.95, suggesting a similar effect for the face classification of adult marmosets in Figure 3D and Appendix A—Table 2. The normalized confusion matrices were compared between the training and validation dataset (Figure 6). It was evident that the automatic face and identity extraction exhibited a high detection accuracy. The background class of the confusion matrix showed frequently predictions as marmoset faces and collar beads for the training (Figure 6A) and validation set (Figure 6B). However, it does not necessarily indicate incorrect predictions or misclassifications. Instead, these values were mostly explained by multiple detections of the same object class. For instance, additional marmoset faces were predicted when multiple animals were present within a single video frame. The long collar structure or motion blur of the marmosets could also cause multiple detections of beads that belong to the same collar. This also corresponded to the high precision and recall scores observed across prediction classes (Figure 5D), suggesting that the increased background false positives were mainly related to the object-count discrepancies, instead of poor detection performance.

**Figure 5.**
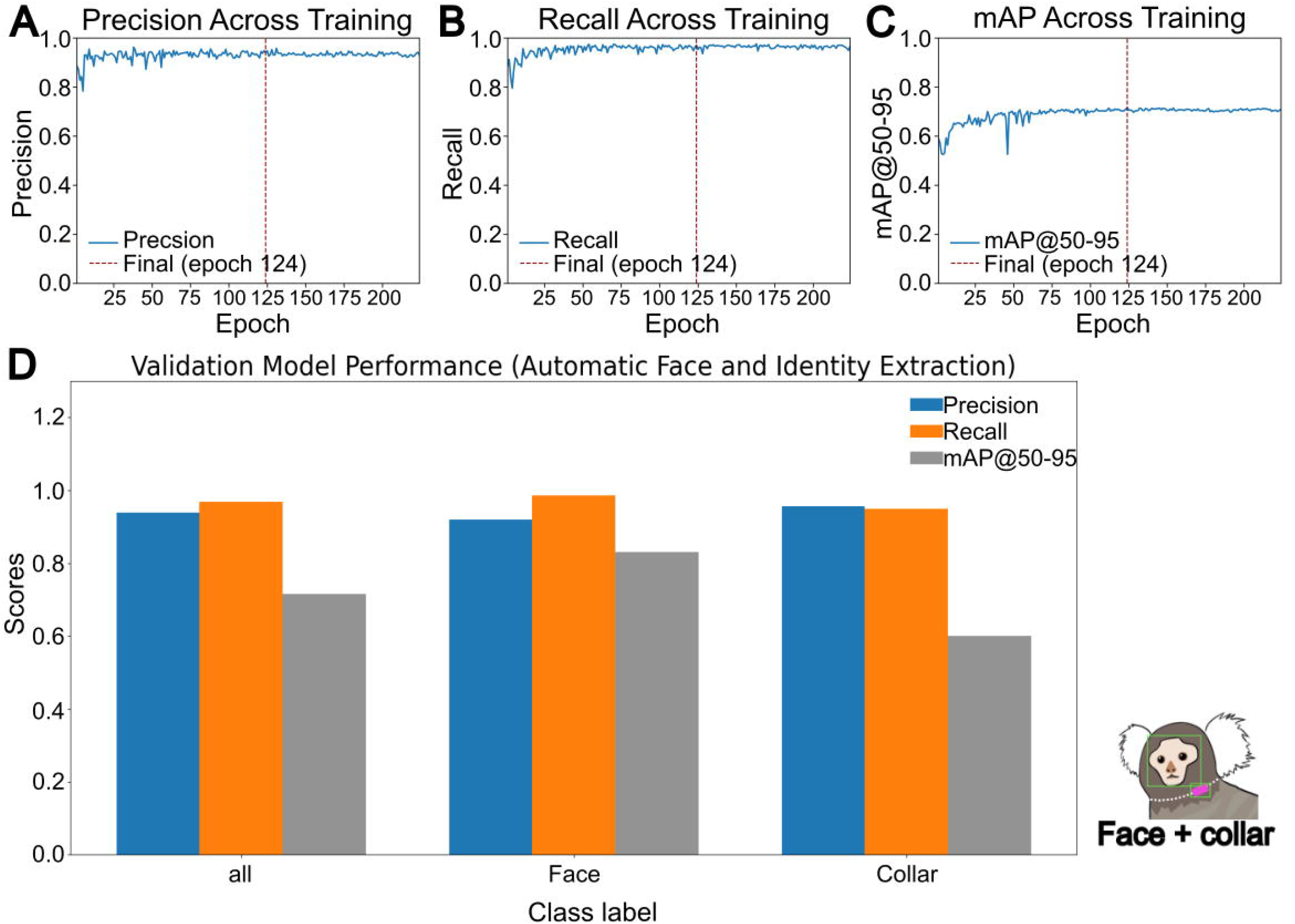
Training performance of the automatic facial and identity extraction model. (A) Precision of all detection classes across training epochs. The red dotted line denoted the final model with the best performance at training epoch 124. (B) Similar to (A), except for overall recall. (C) Similar to (A), except for the mAP at the IoU at 0.5:0.95. (D) Overall Model precision, recall, and the mAP@50-95 (IoU = 0.5:0.95) for each label class.

**Figure 6.**
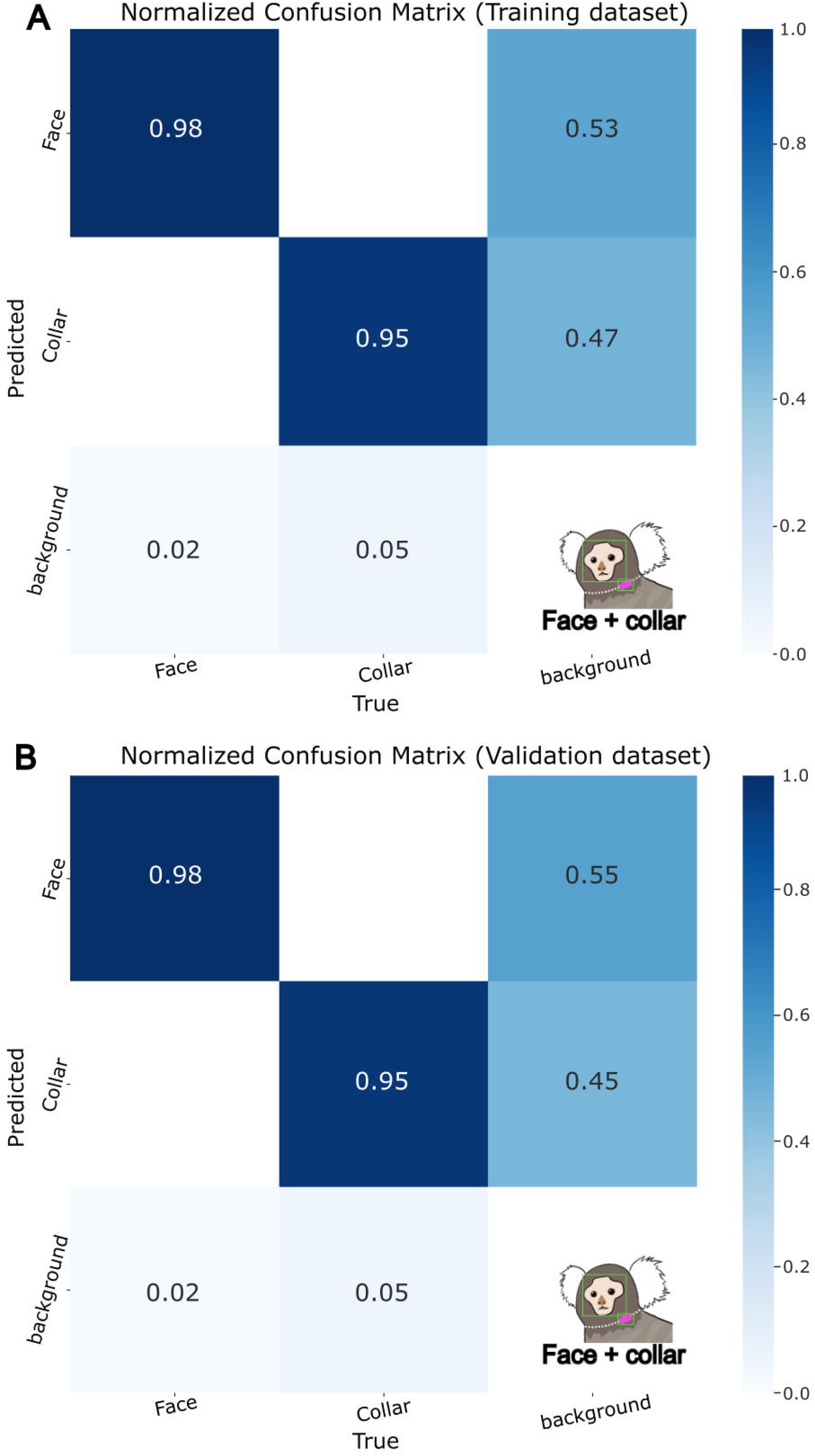
Normalized confusion matrix per-class classification across the 2 label classes in the automatic identity extraction model. Proportion was generated by the (A) training dataset and the (B) validation dataset, showing whether certain classes were frequently mislabelled as a different class.

To test the performance of the automatic face and identity extraction, we used different short clips of new marmosets. We showed that our automatic face and identity extraction pipeline could detect and localize new marmoset faces and collar beads at different ages. Video 2 showed an example of the detection of faces and bead-color labels for two new younger marmosets at 7 months and 17 months, with the confidence threshold of the detection set at 0.6.

### Comparison of identification accuracy in young marmosets across developmental stages

The training and validation datasets of the young marmosets were annotated automatically using the automatic facial and identity extraction. All automatically generated labels were reviewed by the experimenter, and mislabeled images were removed prior to the model training. The training performance curves were presented in Figure 7A – C and Figure 7—figure supplement 1. The final weights were selected at the training epoch of 245, where the training performance metrics achieved a plateau. This program achieved the best performance matrices at this epoch, with precision score at 0.979, recall at 0.975, and mAP@50-95 (IoU = 0.5:0.95) at 0.892. In the young marmoset faces and their collar bead label classes, our face classifier reached high precision and recall performances using the validation dataset (Figure 7D, see Appendix A—Table 3). The precision and recall scores for the label classes were: 1) face of Young1: precision = 1, recall = 0.935; 2) face of Young2: precision = 0.958, recall = 1; 3) collar of Young1: precision = 0.998, recall = 1; 4) collar of Young2: precision = 0.963, recall = 0.964. In terms of localization accuracy, we found a similar pattern of reduced mAP@50-95 (IoU = 0.5:0.95) in bead-color class (mAP@50-95 = 0.808, 0.855), compared to the marmoset faces (mAP@50-95 = 0.95, 0.947). We further computed the normalized confusion matrices of this face classifier using the training and validation dataset. We found that our face classification was relatively accurate across detection of most class labels (Figure 8). The normalized confusion matrices showed high accuracy and consistency of most marmoset faces and collars detection in training (Figure 8A) and validation (Figure 8B) tests, with some exceptions. Particularly, the background was frequently identified as the collar of Young2 marmoset. This elevated background score was likely contributed by the multi-color design of the Young2 marmoset collar, making it more difficult to distinguish compared to collars with a single bead color. In this occasion, if one bead is occluded, blurred, or outside the field of view, the other visible collar bead could affect the prediction and lead to an incorrect identification from the ground truth.

**Figure 7.**
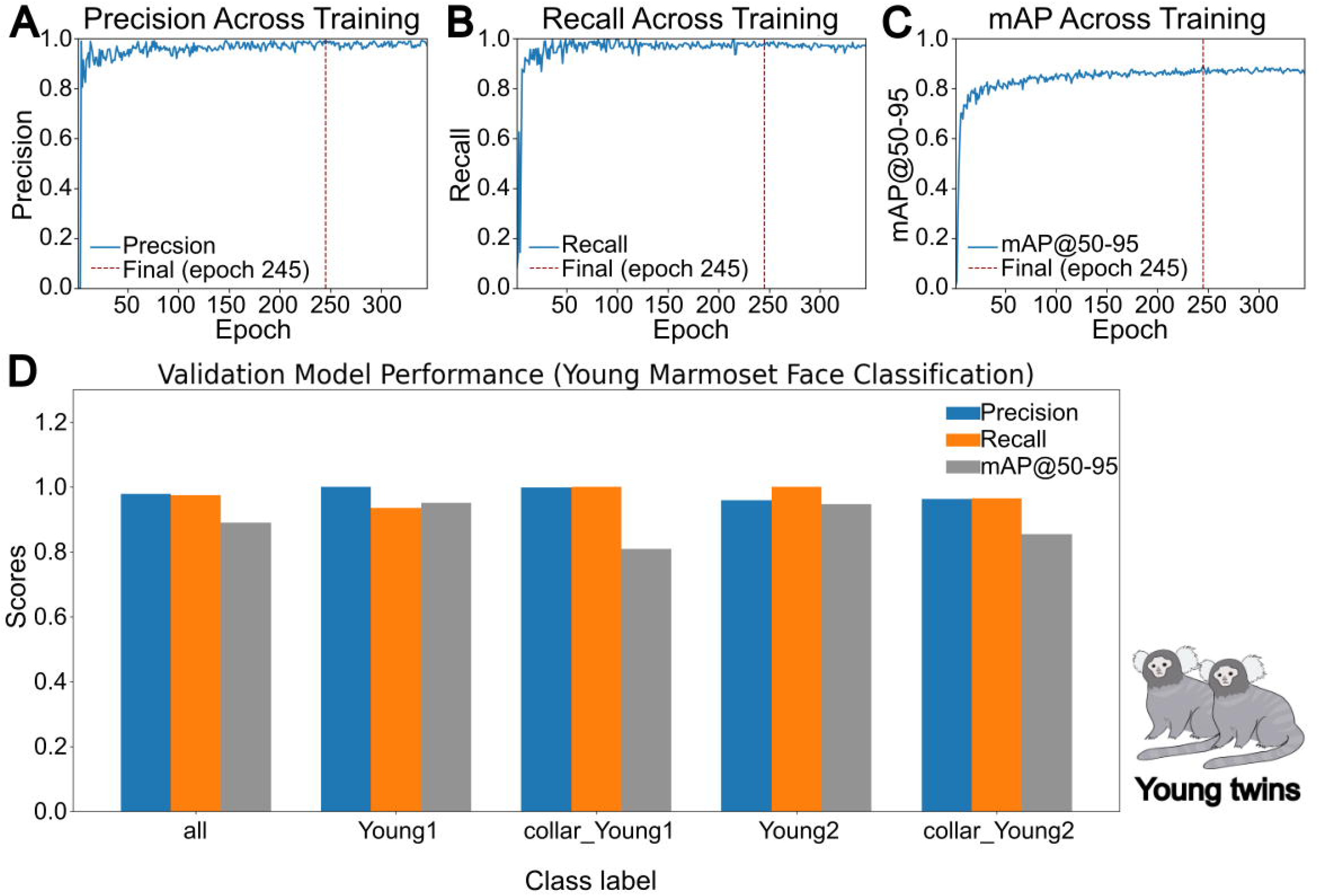
Training performance of the multi-marmoset face recognition model for young marmosets at 7 months. (A) Precision of all detection classes across training epochs. The red dotted line denoted the final model with the best performance at training epoch 245. (B) Similar to (A), except for overall recall. (C) Similar to (A), except for the mAP at the IoU at 0.5:0.95. (D) Overall Model precision, recall, and the mAP (IoU = 0.5:0.95) for each label class.

**Figure 8.**
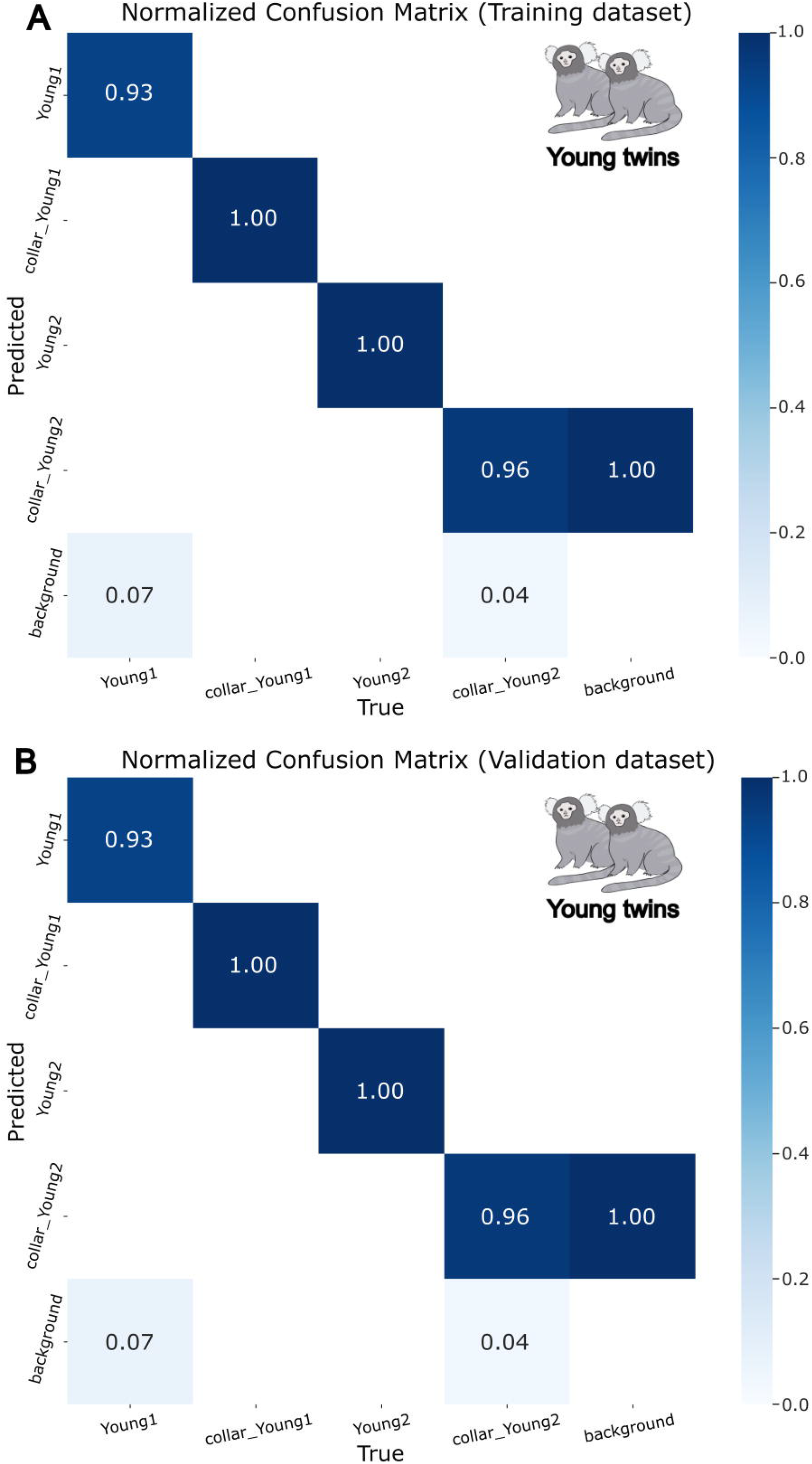
Normalized confusion matrix per-class classification across the 4 label classes in the young marmoset recognition model. Proportion was generated by the (A) training dataset and the (B) validation dataset, showing whether certain classes were frequently mislabelled as a different class.

To test the detection performance of the face classifier trained on 7-month-old young marmosets across different developmental stages, we used short clips of the two young marmosets at the age of 7 months (Video 3, Video 4) and 11 months (Video 5, Video 6). Video 3 - 6 presented the example detections of the young twins at different ages, showing the difficulty of identifying and distinguishing between twin marmosets.

By combining the identification of marmoset faces and collar beads, the program showed valid detection of the correct identity of individual marmosets. However, the detection of the twin marmosets also posed various challenges for computer vision and resulted in mislabeling, including highly similar faces of twin marmosets, face blur or mislabeling due to complex backgrounds and dark lighting conditions (Video 3), fur occluding collar beads, marmosets only showing faces during a short time intervals (Video 6), similar or same color of collar beads between two marmosets (Video 5, Video 6), and fewer training data (young marmosets’ model only included 449 training and validation images for two marmosets at 7 months old).

We tested the prediction performance without collar and its longitudinal application, using the face-only prediction on the young marmosets at 11 months and 16 months (Video 7, Video 7—video supplement 1, Video 7—video supplement 2). The face classifier correctly identified the young twin marmosets solely based on their facial features, indicating that facial identity classification was performed independently of collar information and that the collar beads acted only as an auxiliary cue rather than the main classifier of the system (Video 7).

### Facial similarities of marmosets affect the performance of the facial detection and identification model

We quantified the inter-individual face similarities using cosine similarity and Euclidean distance between mean normalized embeddings of marmoset pairs. We included both adult and young marmosets’ dataset and computed the face similarity based on the 4 relationship pairs: 1) mother-father; 2) father-son; 3) mother-son; and 4) twin-twin. To visualize across the two independently trained model, the inter-individual face similarity scores were normalized using z-score within each model (Figure 9, Figure 9—figure supplement 1 - 2).

**Figure 9.**
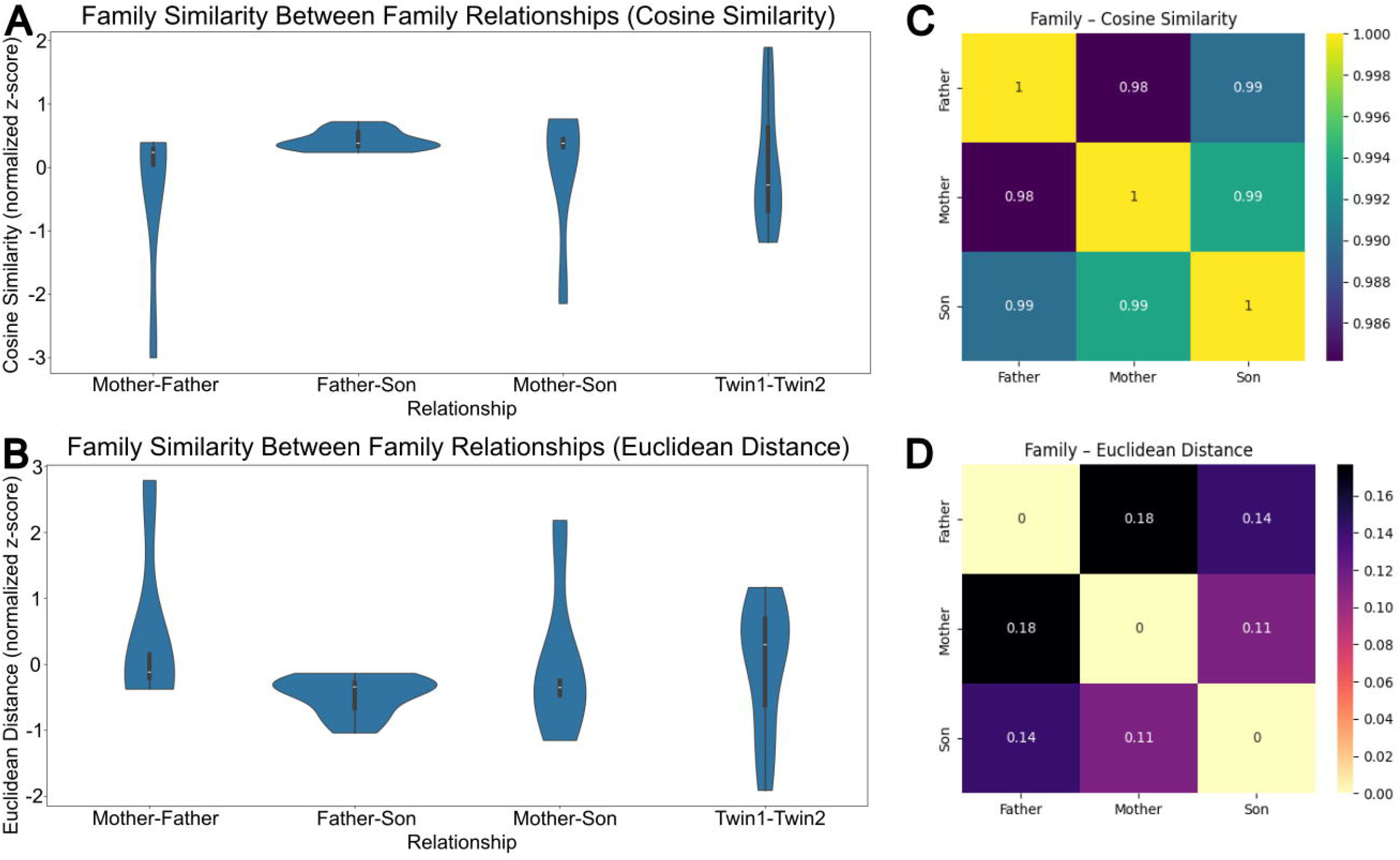
Across-model visualization of the face similarity between marmoset pairs. Four types of family relationships (mother-father, father-son, mother-son, and twin1-twin2) were compared, based on the training results of adult and young marmosets. The similarity was calculated using (A) cosine similarity and the (B) Euclidean distance. Heat map of the (C) cosine similarity and (D) Euclidean distance was plotted between the 3 relationship pairs in the adult family. The cosine similarity score ranged from 0 (very different) and 1 (exactly same) for marmoset faces. The Euclidean distance score ranged from 0 (exactly same) and 1 (very different) for marmoset faces.

Within the model trained using the adult marmoset family, both face similarity measures showed a consistent pattern of across the family relationships. In the adult family, we found that father-son pair showed the highest face similarity values, with the cosine similarity at a positive mean z score (z = 0.43) and the Euclidean distance at a negative mean z score (z = −0.47) (Figure 9C and D). This pattern suggested that father-son pair exhibited a higher-than-average similarity relative to other family relationship. On the other hand, mother-father relationship showed the lowest face similarity in the family (negative cosine similarity z = −0.38, positive Euclidean distance z = 0.43) compared to the adult marmosets’ model mean. The similarity of the mother-son pair was calculated at cosine similarity z = −0.05 and Euclidean distance z = 0.04, indicating an intermediate face similarity in the adult family.

The young marmosets’ identifier was independently trained specifically on the two twin marmosets. Thus, the normalized face similarity z scores were centered near zero and not informative of the twin similarity in family relationship comparison (Figure 9). However, we found a narrow distribution of the image embeddings between the twin marmosets, indicating a nearly invariant facial embeddings and structure between the two twins, with an extremely low raw variance for cosine similarity (std = 0.0005) and Euclidean distance (std = 0.0034).

We performed the statistical tests only on the face classifier for the adult marmoset family, as the twin marmoset model only involved two individuals and thus not valid for within-model statistical analysis (Table 2, 3). Cosine similarity showed a trend of lower similarity in mother-father pair compared to the father-son pair (t = –1.941, p = 0.083 > 0.05, Cohen’s d = –0.868 (large effect)), though not significant. Differences between other relationship pairs were not significant (Table 2). Our analysis on the Euclidean distance indicated a significant difference between mother-father and father-son pairs (t = 2.28, p = 0.046 < 0.05, d = 1.02 (large effect)), while the remaining relationship pairs appeared non-significant (Table 3).

**Table 2.** Statistical analysis of inter-individual face similarity (cosine similarity) between different family relationships, within the adult marmoset family. Comparison of t-statistics, p-value, and Cohen’s d on cosine similarities were compared between the 3 family relationships. The relationship pairs that showed significant differences were bolded.

| Relationships | t-value | p-value | Cohen's d |
| --- | --- | --- | --- |
| Mother-father vs. Father-son | -1.941 | 0.083 | -0.868 |
| Mother-father vs. Mother-son | -0.613 | 0.548 | -0.274 |
| Father-son vs. Mother-son | 1.460 | 0.177 | 0.653 |

**Table 3.**
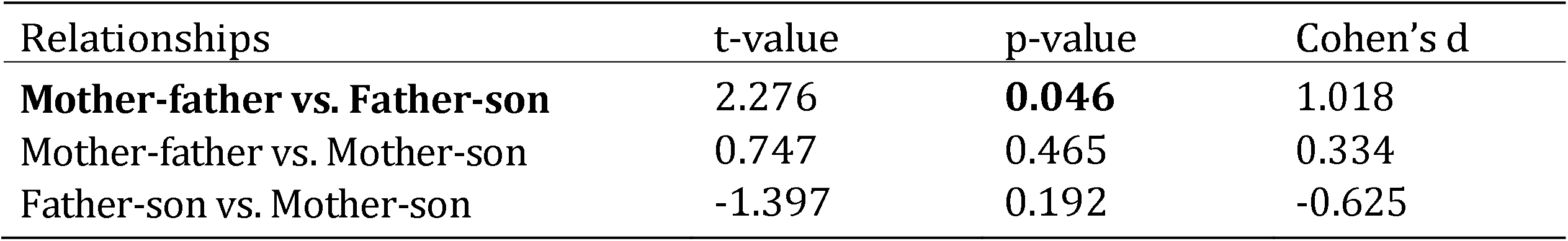
Statistical analysis of inter-individual face similarity (Euclidean distance) between different family relationships, within the adult marmoset family. Comparison of t-statistics, p-value, and Cohen’s d on Euclidean distances were compared between the 3 family relationships. The relationship pairs that showed significant differences were bolded.

### A real-time marmoset identification program, based on the trained networks

We developed a real-time interface for detecting and identifying the marmosets, based on the trained models of multi-marmoset classification. The real-time footages were acquired using the same camera and experimental setup as the training videos (see examples in Video 1 – 6). Our real-time program processes the real-time footage like the detection of the offline collected videos, using the best model selected from each multi-marmoset face classification training model. Prior to running the real-time program, the experimenter first trains the multi-marmoset face classification model based on the specific subjects of interest. Next, the model with the best performance from training are selected and inputted into the real-time program. No programming is required from the experimenter using this real-time marmoset face identification program. The experimenter clicks a run button on the program console to initiate the real-time detection. While this program is running, the experimenter can check the frame-by-frame label detection results (30 frames per second) displayed on the screen. Potential mislabeling are mitigated by combining the marmoset faces and collar beads detection, with greater weights assigned to the collar bead detections than the marmoset face (example mislabelling in Video 5 and Video 6). The program automatically records the most frequently detected marmoset identity across 30 frames (approximately one second). At the end of the experiment, the program is terminated by clicking a stop button of the python console. After the completion of the experiment, the full record of marmoset identity detection results is available for review by the experimenter.

## Discussion

We developed a real-time computer vision program for automatic marmoset identity recognition, using their facial features and uniquely color-coded bead collars. By combining the automatic annotation of the real-time footage with the individual-specific classification (i.e. faces and collar beads), our program allows continuous identity tracking during behavioral experiments, with minimal human interference. Utilizing the object detection YOLOv8 model based on pre-trained image networks, we adapted it to a non-human primate (i.e. marmosets) face dataset and applied it in real-time tracking. Pose estimation tools have been widely used in characterizing animal behavior; however, these tools require extensive computational power thus limiting the identification between visually similar individuals, especially animals housed in family unit (Camilleri et al., 2023; Gill et al., 2025; Lauer et al., 2022; Mathis et al., 2018; Nath et al., 2019). Notably, the main goal of the current system is marmoset identity recognition, instead of pose estimation or behavioral analysis, such that identity characterization is not dependent on posture cues or movement tracking. Thus, we selected the YOLOv8 object detection algorithms for the development of our real-time marmoset face recognition pipeline. Among the YOLOv8 pre-trained models, we selected the light-weighted YOLOv8 nano model, which provided the optimal balance between detection accuracy and computational inference speed, supporting its feasibility for the final real-time marmoset identity recognition program.

Our real-time marmoset recognition pipeline includes two face classification for both adult and young marmoset, with an additional automatic face and identity extraction program for all marmosets. The models presented in this paper achieved reliable detection accuracy across adult marmosets’ and twin marmosets’ datasets, with detection accuracy improving with increased amount of training images. While comparing between the adult and young marmoset datasets, we found that the adult marmosets’ face classifier, trained with a larger number of varied images, showed more reliability and efficiency in marmoset identity recognition. Moreover, we anticipate that experimenters can integrate this program into behavioral experiments, as an automated marmoset identity extraction and classification tool for real-time video monitoring. Once fully trained on recognizing subject marmosets, this pipeline operates automatically and can be applied across individuals of different ages, with minimal manual work needed. While existing marmoset identification approaches usually utilize visible markers, Radio Frequency Identification (RFID), or observation, the manual works and human interventions involved can impact animal behaviors, especially during their behavioral task performance. The facial identification tool aims to collect data from marmosets without having experimenters to check the identity continuously, instead of outperforming the experimenters’ role. Its automated pipeline substantially reduces the time and work required for traditional manual identity labeling, while maintaining an expert-level human performance and reproducibility across experimenters (95.83% average accuracy for animal health technicians, responsible for daily health check and husbandry while lab experiments ranges between 25 to 80% of accuracy depending on the amount of time spent with each animal, Supplementary figure 1). The tool’s advantages are particularly efficient for large datasets and longitudinal studies, where manual identity labeling becomes difficult, as variability and errors increase along with dataset size and experimenter number.

The motivation for this real-time marmoset identity recognition program was to develop an easy-to-use, generalizable pipeline that could be applied across different marmosets and lab environments, such as using larger field of view or phone cameras (Figure 10). The pipeline was designed to have no specific hardware requirements and can be implemented for any standard recording device, including any commonly available cameras, primate chair system, and computer-based device. Based on the collected individual marmoset face data, automatic extraction program allows consistent identity annotation of marmoset faces and collar beads, ensuring accurate and stable identity interpretation during the marmoset face classification model training. The experimenter only needs to run the Python pipeline scripts to perform the real-time marmoset identification during experiments, without the need to manually label large marmoset identity datasets. Moreover, the system operates directly based on the raw video input from the camera, with no pre-processing such as video cropping or resolution modification required. The identity recognition system recognizes each marmoset based on pre-defined labels from collected videos of individuals, without other external tools such as RFID systems (Pereira et al., 2023). We focused the marmoset identification on individual-specific facial features and uniquely colored collar beads to ensure the pipeline robustness across various recording conditions, regardless of lighting condition, camera angle, or animal posture. Together, this pipeline design and setup enable fast, reliable, and accurate identity recognition for an efficient real-time monitoring of multiple animals in complex experimental environments. However, with the high visual similarity between closely related marmoset family members, the facial features and the collar beads must be carefully integrated in our pipeline design, as even experienced human observers can misidentify marmoset twins.

**Figure 10.**
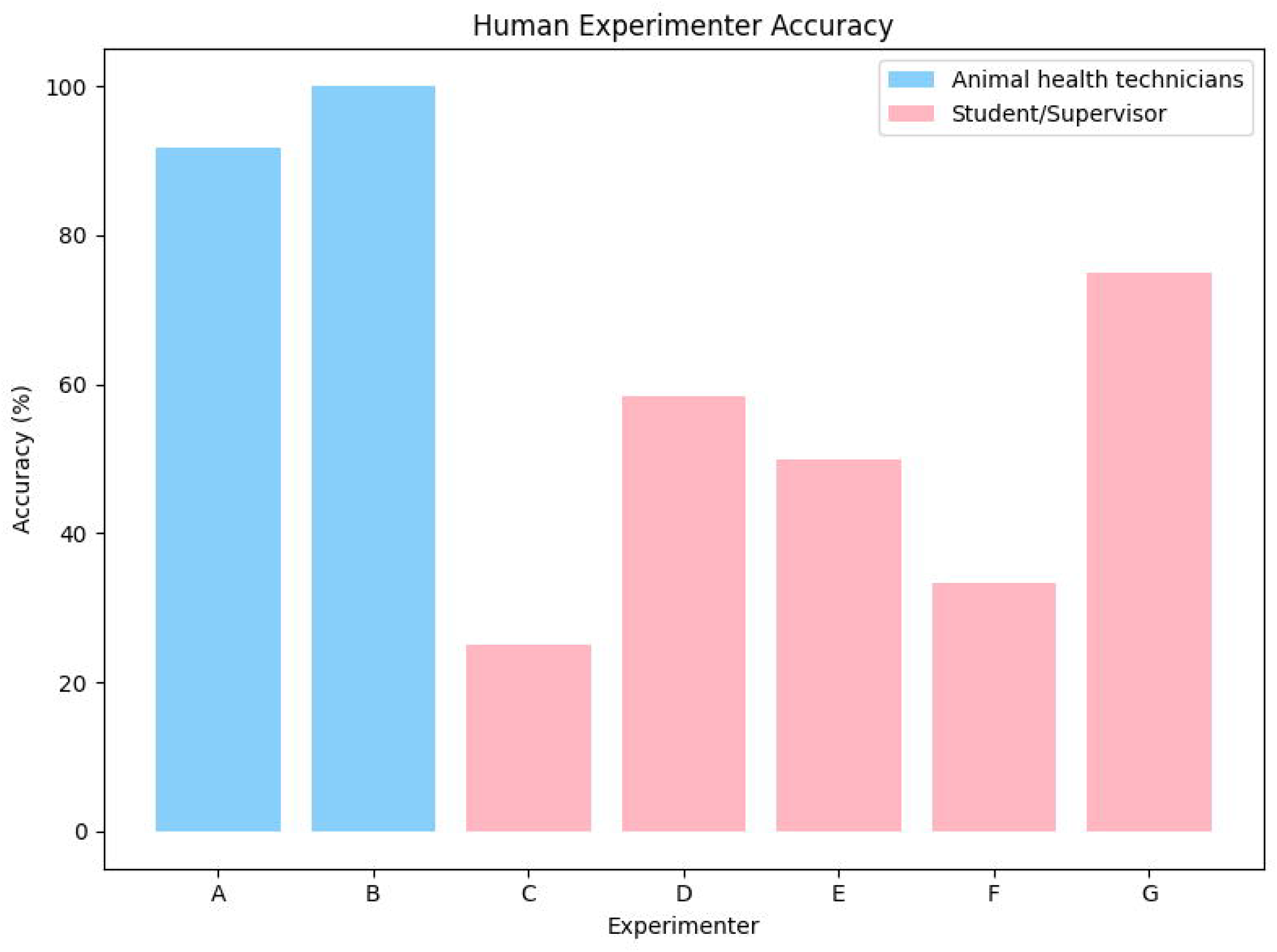
Examples of marmoset face detection and classifier for in-cage and open-field marmoset images. This image illustrated the (A, B, D, E) detection of marmosets faces and (C) their identities across different cameras, environments, and marmoset behaviors. Additionally, it included scenarios where animals were housed in-cage (A – C) or moving freely (D – E). The open-field marmoset videos (D - E) were obtained from the Youtube Billabong Zoo, Koala & Wildlife Park: https://www.youtube.com/watch?v=lDJNk4rjqqQ.

Because of the high extent of natural facial similarity between related marmosets, our real-time pipeline can experience challenges in identifying closely related individuals, as the results in Figure 7 - 9, and Video 3 - 6 indicate. Fur textures, coloring patterns, and genetic relatedness contribute to the morphological similarities, making it challenging for human observers when identifying individual primates (Alvergne et al., 2009; Guan et al., 2023; Leopold and Rhodes, 2010). Facial features serve as the main and intrinsic biometric identifier for each marmoset, providing a unique source of individual recognition. Since collar-based confirmation could be affected by visibility limitations, we implemented the uniquely color-coded bead collars as an auxiliary cue to provide additional confirmation in identity prediction. For example, this issue can be caused by identical or similar bead colors between individuals (Video 5, 6) and beads that are occluded by fur (Video 3 - 6). In addition, collar beads may change over time or not be worn by all animals. An alternate experimental solution is to improve collar visibility, including using distinct color code across individuals within a family, increasing the size of the beads, or increasing collar beads number to reduce occlusion. Moreover, the performance of the system is strongly dependent on the amount and variability of the training data, with identity classification improving as more marmoset images are involved in the model training. This relation is specifically important when characterizing between young and adult marmosets, as facial features may change across marmosets’ developmental stages. Images from both the juvenile and adult periods of the same marmoset can be included in the model training, which could improve model performance and generalizability across developmental stages, and avoid repeated training of the individual recognition model at different ages.

Because our pipeline was designed specifically for marmoset identity recognition, it was optimized for characterizing individuals within a defined housing unit, which usually represents as a family or pair group. With one separated model trained per family unit, our system can utilize distinct collar colors as an additional identifier when available, while facial features performed as the main biometric marker. Even though multiple marmosets with visually similar faces may present close to the camera, the additional collar information can improve confidence in identity prediction without replacing facial recognition as the primary mechanism of identification (Video 7). Moreover, while changes in the recording environment setup (e.g. lighting condition, camera angles, etc.) does not affect the model performance, the model needs to be adjusted when family members or composition changes. In particular, the introduction of new members (e.g. newborns) requires assignment of new colored collar beads and collection of face images, thus it is necessary that the recognition model of this marmoset family is retrained.

This system may be particularly useful when experimenters need to know which individual is performing a specific task for cognitive and behavioral experiments (Kangas et al., 2016; Kangas and Bergman, 2017; Marshall and Ridley, 2003). In these experiments, especially those involving long-term or continuous behavioral responses, experimenters often need to be present to record animal identity or to manually review video recordings after the experiment. This may distract the animals from their task and affect their behaviors, while also demanding sustained attention and large time commitment for the experimenters. Our pipeline resolves this limitation by automatically detecting and recording marmoset identity throughout the experiment. A typical system requires the experimenter to only collect video clips from each marmoset, verify the automatically annotated marmoset faces and collar beads, and initiating the training of a family-specific face classification. This process takes approximately 20 - 30 hours of time. Once trained, the system operates automatically to collect real-time identity and can work to present subject-specific behavioral or cognitive tasks based on the identity of the detected animal, with no work or presence needed on the user’s end. This tool has already been implemented in touchscreen-based marmoset cognitive tasks, including pairwise visual discrimination paradigm.

Future extensions can further improve the detection accuracy and the pipeline utility. Firstly, the pipeline can be easily combined with additional programs to characterize the detailed facial features of the marmosets (Correia-Caeiro et al., 2022; Kawaguchi et al., 2023). Considering differences in their eye distance, mouth shape, and fur coloring pattern, in addition to the global facial structure, we can improve the identity prediction, particularly between two closely related marmosets with similar faces. Also, aside of the colored collar beads, experimenters can label marmosets’ identities using other visual markers, including color dyes on marmoset ear tufts. Using a more evident visual marker could be helpful for accurate identity prediction and avoid mislabeling. Moreover, it would be possible to integrate pose estimation tools, such as DeepLabCut and MarmoPose (Cheng et al., 2025; Lauer et al., 2022), with this marmoset recognition pipeline, thus allowing the estimation of the postures and behaviors of specific animals of interest. While we developed this pipeline and demonstrate its utility for common marmosets in laboratory captivity, there is no application restriction of this system. With appropriate training data and experimental design, our pipeline can be applied to other non-human primates in various settings such as lab housing, conservation fields, or even in the wild.

## Data Availability

All code in this paper is publicly available at Github: https://github.com/Jy-Yang-bot/real-time-marmoset-recognition. This repository contains the scripts for pre-processing the images, the main facial recognition pipeline, and the analytic tools. The program generalizability was tested on additional video obtained from publicly available online sources, including a YouTube video “Common Marmoset”, uploaded by Billabong Zoo, Koala & Wildlife Park on 2 June 2020 (https://www.youtube.com/watch?v=lDJNk4rjqqQ).

## Acknowledgments

We would like to thank our animal health technicians D. Hau-Aquino, V. Comtois, C. Hunt; and veterinarians F. Chaurand, and J. Hutta for animal health care and support. We are thankful to M. Gacoin and T. Cook for helpful discussions and feedback on the visualization and format of the manuscript. We thank J. Smith and C. O’Hare-Freire for the design and construction of transparent protective case for the camera. We acknowledge the support of the Government of Canada’s New Frontiers in Research Fund (NFRF), [NFRFT-2022-00051] and by the Fonds de Recherche du Québec–Santé (FRQS), [#347426 and #358082]. Ces travaux ont bénéficié d’un octroi des fonds Nouvelles frontières en recherche du gouvernement du Canada [NFRFT-2022-00051] et du Fonds de recherche du Québec-Santé [FRQS, #347426 et #358082].

## Article and author Information

### Jiayue Yang

- The Neuro, Department of Neurology and Neurosurgery, McGill University, Montreal, Quebec. Canada.
- Integrated Program in Neuroscience, McGill University, Montreal, Quebec, Canada.

#### Contributions

Conceptualization, pipeline development, data collection and curation, dataset labeling, formal analysis, investigation, visualization, methodology, writing – original draft, and writing – review and editing.

### James Wang

- The Neuro, Department of Neurology and Neurosurgery, McGill University, Montreal, Quebec. Canada.
- Integrated Program in Neuroscience, McGill University, Montreal, Quebec, Canada. Contributions: Data collection and curation, writing - review and editing

### Justine Cléry

- The Neuro, Department of Neurology and Neurosurgery, McGill University, Montreal, Quebec. Canada.
- McConnell Brain Imaging Centre, The Neuro, Montreal Neurological Institute and Hospital, McGill University, Montreal, Quebec, Canada.
- Azrieli Centre for Autism Research, The Neuro, Montreal, Quebec, Canada.

#### Contributions

Conceptualization, methodology, writing – review and editing, project administration, supervision, resources, funding acquisition.

## Figures, Videos, and Tables

### Figures

**Figure S1.**
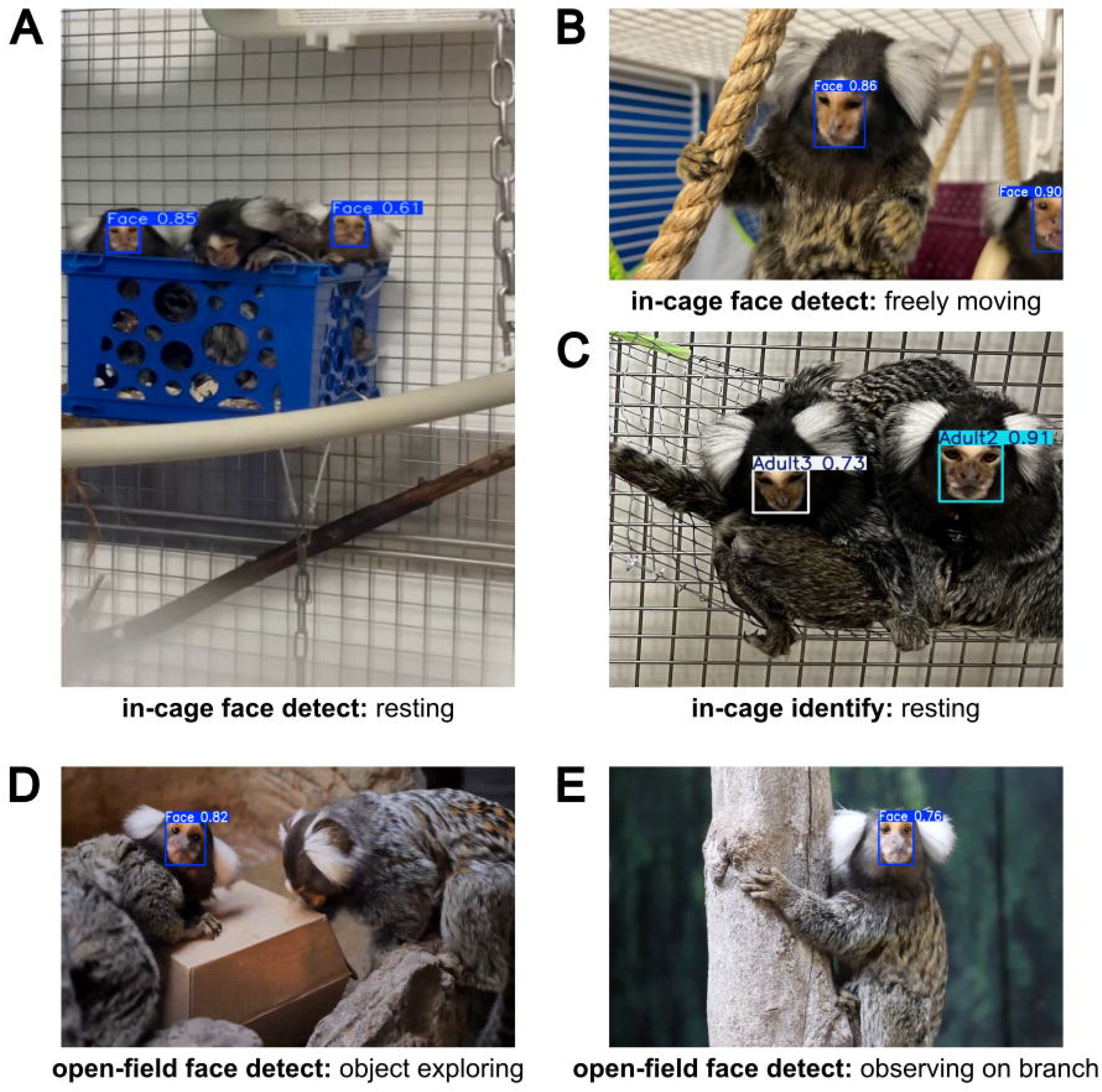
Identification performance of human experimenter across the 5 marmosets. Identification accuracy of 7 human experimenters across the 5 marmosets. The legend denoted the performance of the animal health technicians (blue) and the student/supervisor (pink).

**Figure 3—figure supplement 1.**
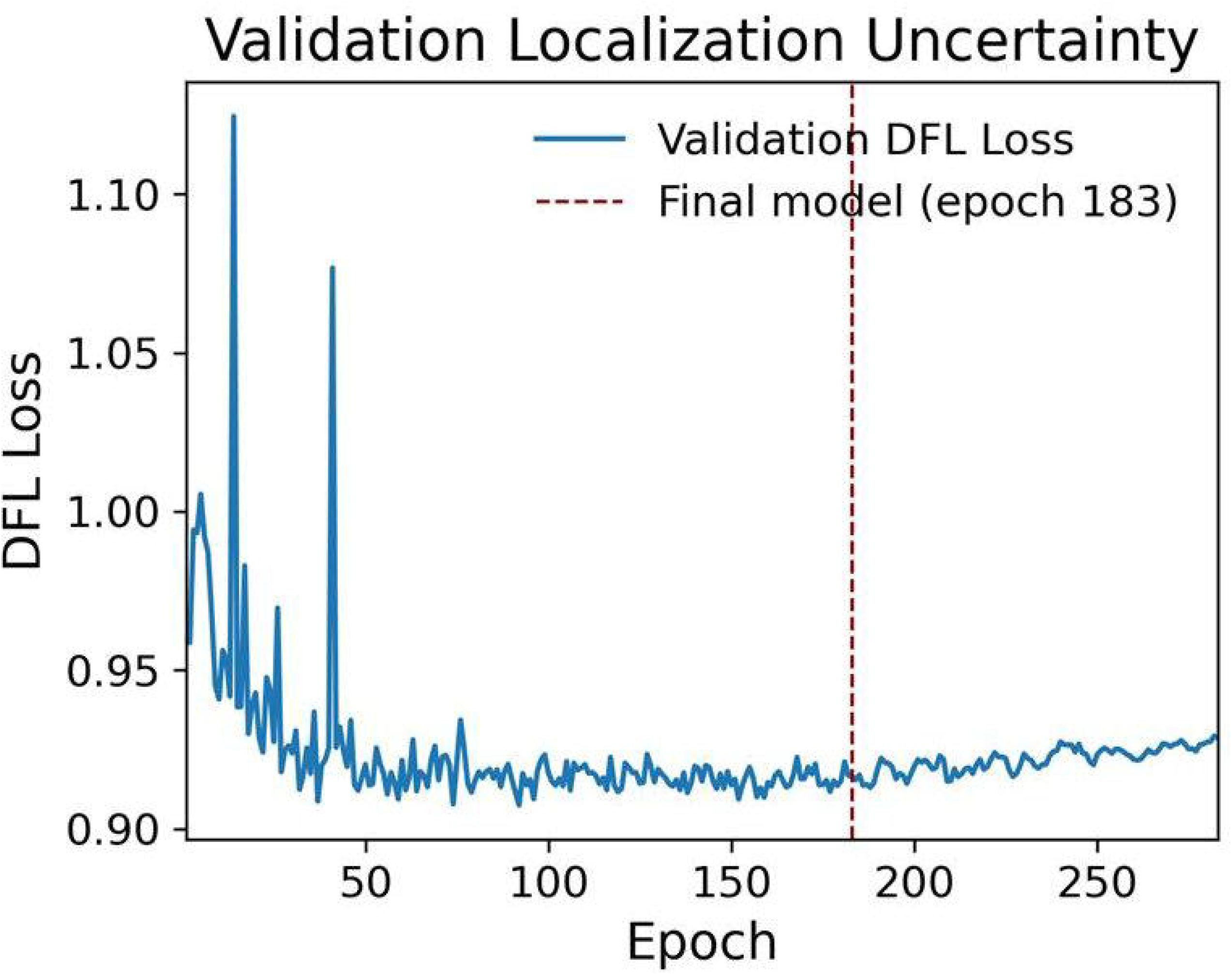
Training performance of the multi-marmoset face classification model for three adult marmosets. The Validation Distribution Focal Loss (DFL) of all detection classes across training epochs. The red dotted line denoted the final model with the best performance at training epoch 183.

**Figure 5—figure supplement 1.**
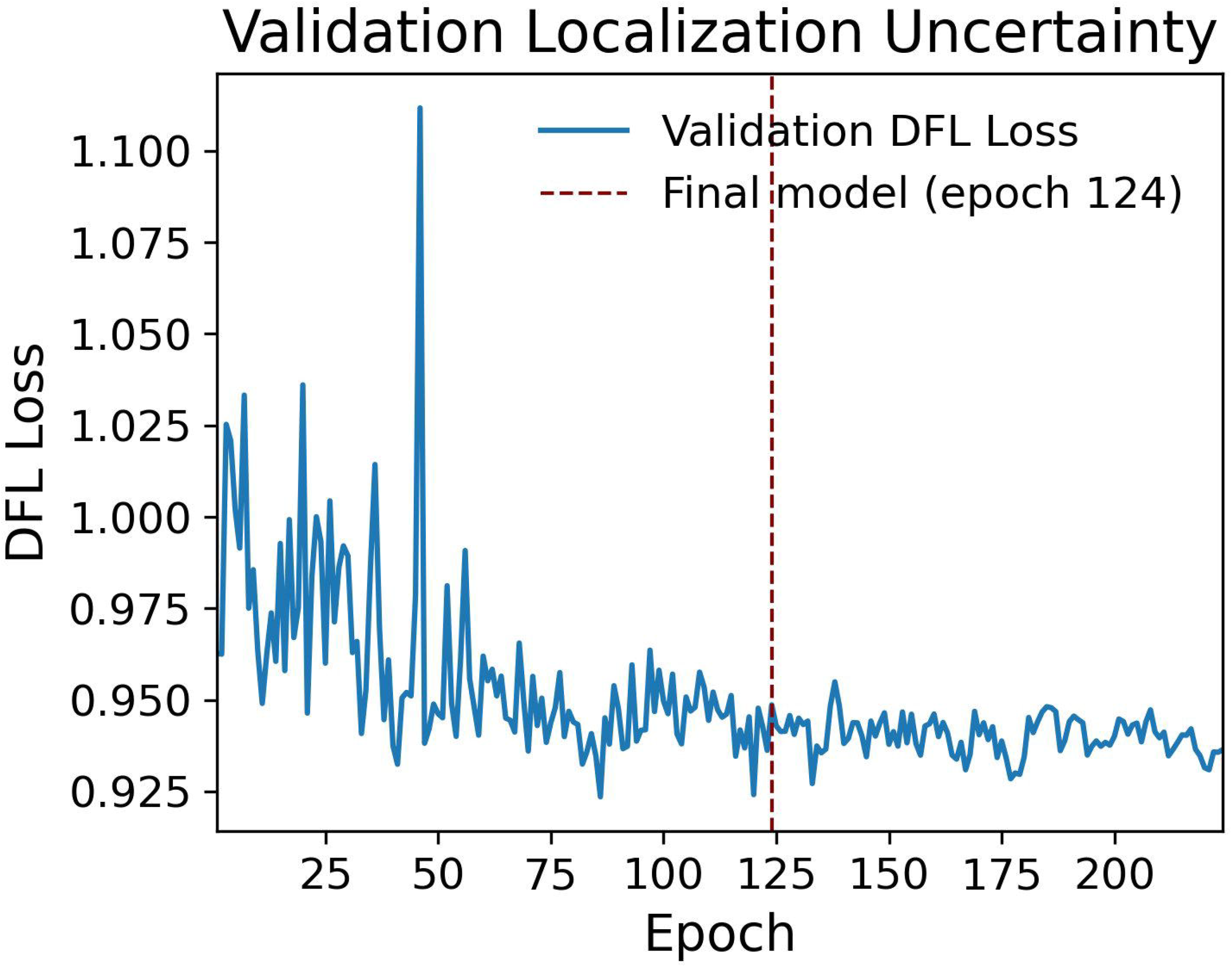
Training performance of the automatic facial and identity extraction model. The Validation Distribution Focal Loss (DFL) of all detection classes across training epochs. The red dotted line denoted the final model with the best performance at training epoch 124.

**Figure 7—figure supplement 1.**
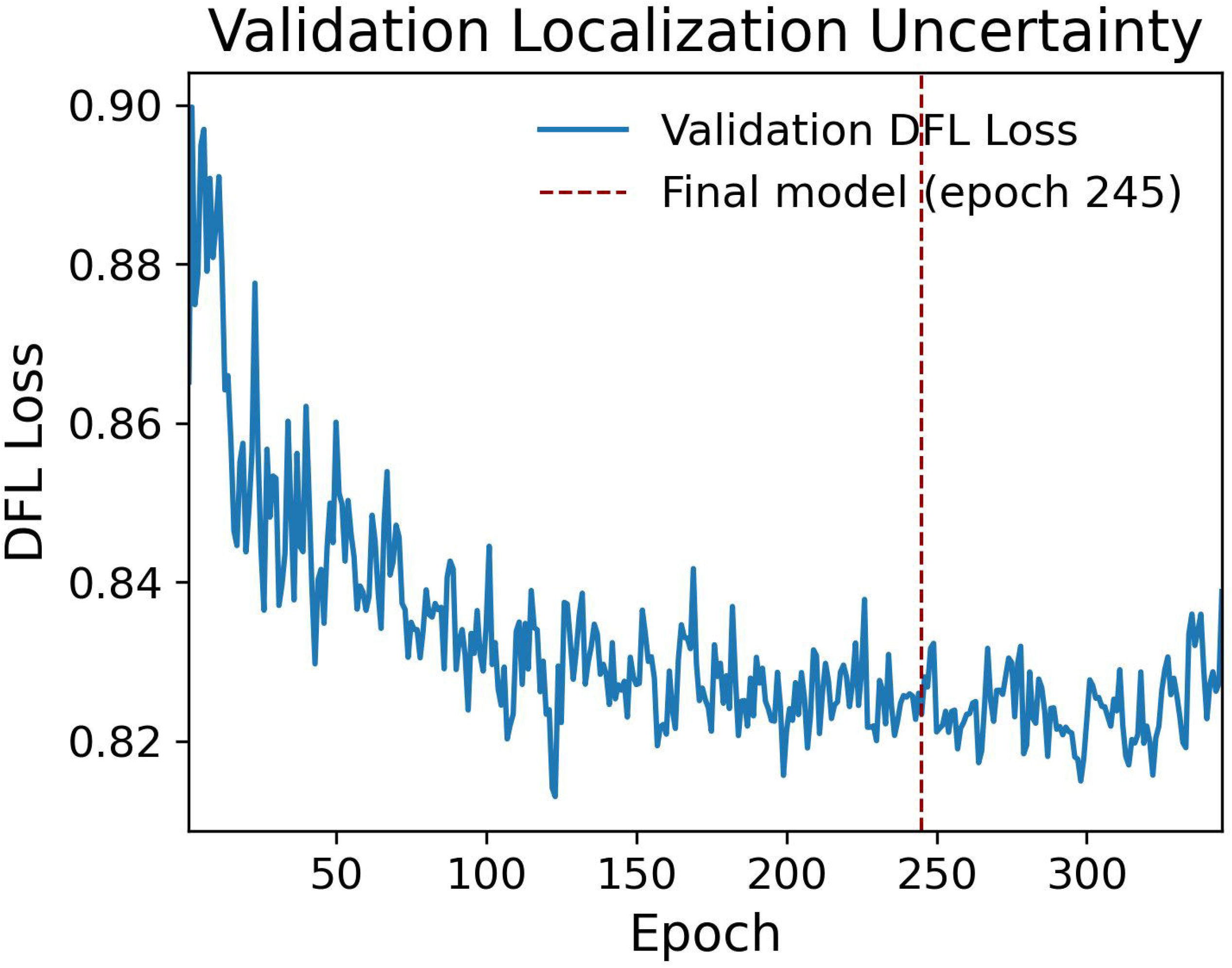
Training performance of the multi-marmoset face classification model for two young marmosets. The Validation Distribution Focal Loss (DFL) of all detection classes across training epochs. The red dotted line denoted the final model with the best performance at training epoch 245.

**Figure 9—figure supplement 1.**
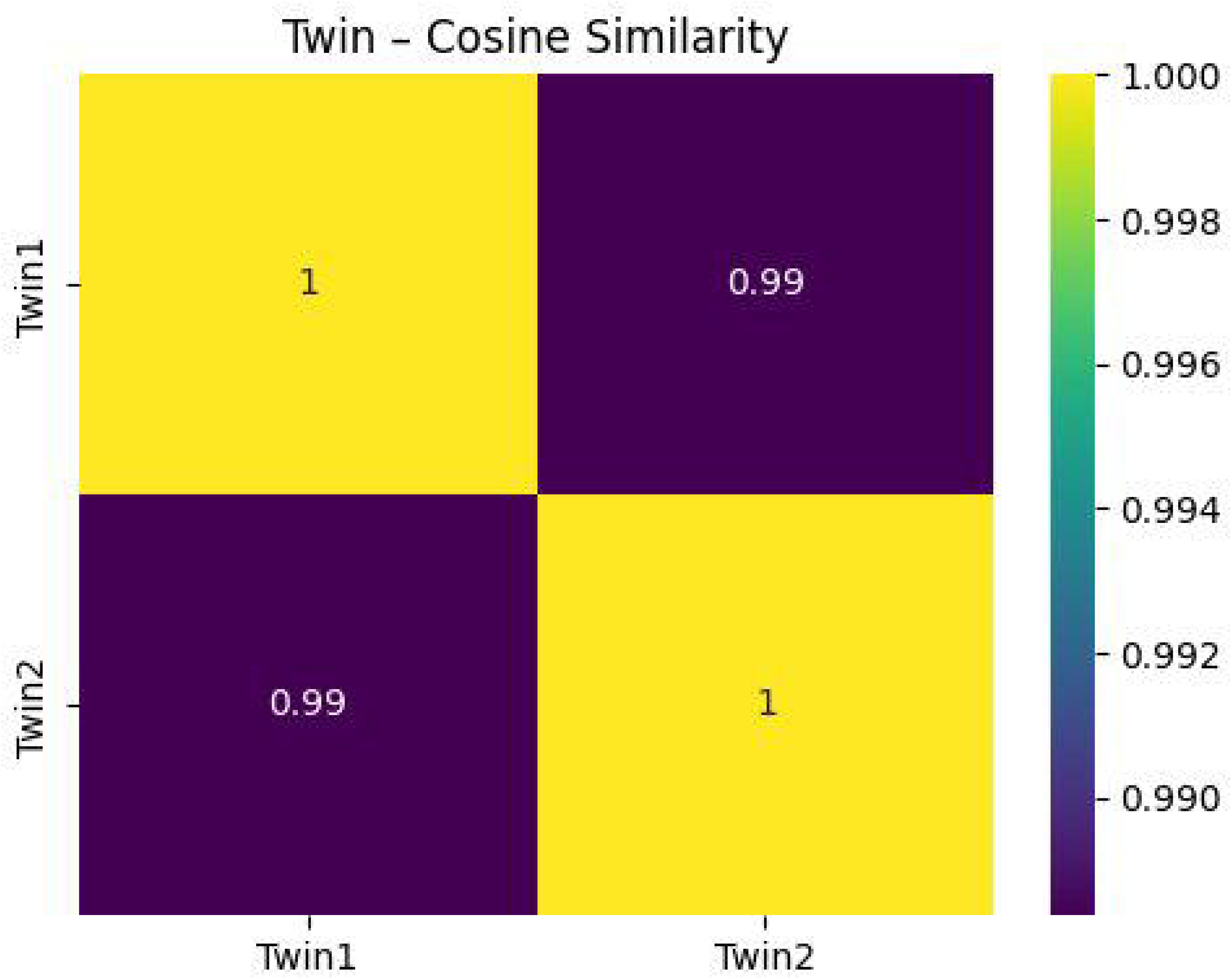
Heat map of the cosine similarity between marmoset faces in the young family. The two young marmosets (twins) were represented by their family, with the similarity score calculated between the two twins. The cosine similarity score ranged from 0 (very different) and 1 (exactly same) for marmoset faces.

**Figure 9—figure supplement 2.**
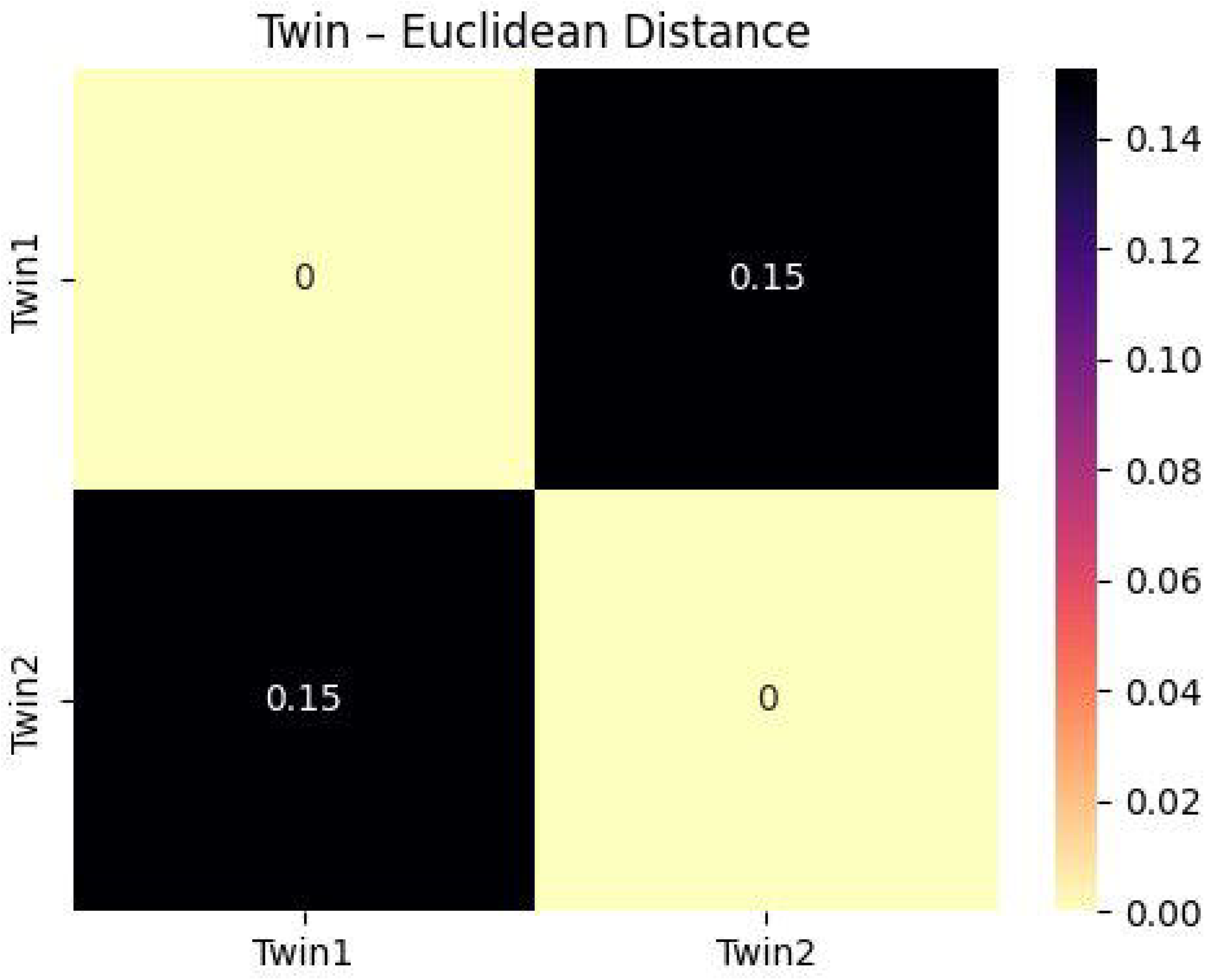
Heat map of the Euclidean distance between marmoset faces in the young family. The two young marmosets (twins) were represented by their family, with the similarity score calculated between the two twins. The Euclidean distance score ranged from 0 (exactly same) and 1 (very different) for marmoset faces.

### Videos

**Video 1. Example annotations of the multi-marmoset face classification from three adult marmosets, from a test video excluded from model training and validation**. Top panel: annotated output video of the classifier. Bottom panel: timeline plot of the detected label classes of adult marmoset faces and corresponding collar beads.

**Video 2. Example annotations of the automatic face and identity extraction from two new marmosets at different age**. Top panel: annotated output video of the extraction model. Bottom panel: timeline plot of the detected label classes of marmoset faces (blue) and collar (cyan) beads.

**Video 3. Example annotations of the multi-marmoset face classification for Young1 marmoset at 7 months**. Top panel: annotated output video of the classifier. Bottom panel: timeline plot of the detected label classes of marmoset Young1 face (blue), Young1 collar beads (cyan), Young2 (pink), and Young2 collar beads (lime green).

**Video 4. Example annotations of the multi-marmoset face classification for Young2 marmoset at 7 months**. Top panel: annotated output video of the classifier. Bottom panel: timeline plot of the detected label classes of marmoset Young1 face (blue), Young1 collar beads (cyan), Young2 (pink), and Young2 collar beads (lime green).

**Video 5. Example annotations of the multi-marmoset face classification for Young1 marmoset at 11 months**. Top panel: annotated output video of the classifier. Bottom panel: timeline plot of the detected label classes of marmoset Young1 face (blue), Young1 collar beads (cyan), Young2 (pink), and Young2 collar beads (lime green).

**Video 6. Example annotations of the multi-marmoset face classification for Young2 marmoset at 11 months**. Top panel: annotated output video of the classifier. Bottom panel: timeline plot of the detected label classes of marmoset Young1 face (blue), Young1 collar beads (cyan), Young2 (pink), and Young2 collar beads (lime green).

**Video 7. Face-only annotations of the multi-marmoset face classification for Young1 marmoset at 16 months**. Top panel: annotated output video of the classifier. Bottom panel: timeline plot of the detected label classes of marmoset Young1 face (blue) and Young2 (pink). The predictions of collar beads were excluded.

**Video 7—video supplement 1. Face-only annotations of the multi-marmoset face classification for Young1 marmoset at 11 months**. Top panel: annotated output video of the classifier. Bottom panel: timeline plot of the detected label classes of marmoset Young1 face (blue) and Young2 (pink). The predictions of collar beads were excluded.

**Video 7—video supplement 2. Face-only annotations of the multi-marmoset face classification for Young2 marmoset at 11 months**. Top panel: annotated output video of the classifier. Bottom panel: timeline plot of the detected label classes of marmoset Young1 face (blue) and Young2 (pink). The predictions of collar beads were excluded.

## Appendix A

**Appendix A—Table 1.** Model performance evaluation of the validation dataset in the adult marmoset recognition model. Comparison of precision, recall, mAP@50, and mAP@50-95 were computed for all the detected label classes in this model.

| Label classes | Precision | Recall | mAP50 | mAP@50-95 |
| --- | --- | --- | --- | --- |
| all | 0.936 | 0.955 | 0.980 | 0.713 |
| Adult1 | 0.829 | 0.945 | 0.957 | 0.841 |
| Adult2 | 0.926 | 0.983 | 0.991 | 0.787 |
| Adult3 | 0.951 | 0.998 | 0.992 | 0.868 |
| collar_Adult1 | 0.968 | 0.966 | 0.988 | 0.603 |
| collar_Adult2 | 0.978 | 0.972 | 0.990 | 0.665 |
| collar_Adult3 | 0.962 | 0.866 | 0.959 | 0.515 |

**Appendix A—Table 2.** Model performance evaluation of the validation dataset in the automatic face and identity extraction model. Comparison of precision, recall, mAP@50, and mAP@50-95 were computed for all the detected label classes in this model.

| Label classes | Precision | Recall | mAP50 | mAP@50-95 |
| --- | --- | --- | --- | --- |
| all | 0.938 | 0.968 | 0.984 | 0.715 |
| Face | 0.919 | 0.986 | 0.990 | 0.831 |
| Collar | 0.957 | 0.950 | 0.978 | 0.600 |

**Appendix A—Table 3.** Model performance evaluation of the validation dataset in the young marmoset recognition model. Comparison of precision, recall, mAP@50, and mAP@50-95 were computed for all the detected label classes in this model.

| Label classes | Precision | Recall | mAP50 | mAP@50-95 |
| --- | --- | --- | --- | --- |
| all | 0.979 | 0.975 | 0.994 | 0.890 |
| Young1 | 1.000 | 0.935 | 0.993 | 0.950 |
| collar_Young1 | 0.998 | 1.000 | 0.995 | 0.808 |
| Young2 | 0.958 | 1.000 | 0.995 | 0.947 |
| collar_Young2 | 0.963 | 0.964 | 0.993 | 0.855 |

